# diffONT: predicting methylation-specific PCR biomarkers based on nanopore sequencing data for clinical application

**DOI:** 10.1101/2025.02.17.638597

**Authors:** Daria Meyer, Emanuel Barth, Laura Wiehle, Manja Marz

## Abstract

DNA methylation is known to act as biomarker applicable for clinical diagnostics, especially in cancer detection. Methylation-specific PCR (MSP) is a widely used approach to screen patient samples fast and efficiently for differential methylation. During MSP, methylated regions are selectively amplified with specific primers. With nanopore sequencing, knowledge about DNA methylation is generated during direct DNA sequencing, without any need for pretreatment of the DNA. Multiple methods, mainly developed for whole-genome bisulfite sequencing (WGBS) data, exist to predict differentially methylated regions (DMRs) in the genome. However, the predicted DMRs are often very large, and not sufficiently discriminating to generate meaningful results in MSP creating a gap between theoretical cancer marker research and practical application, as no tool currently provides methylation difference predictions tailored for PCR-based diagnostics. Here we present `diffONT`, which predicts differentially methylated primer regions, directly suitable for MSP primer design and thus allowing a direct translation into practical approaches. `diffONT` takes into account (i) the specific length of primer and amplicon regions, (ii) the fact that one condition should be unmethylated, and (iii) a minimal required amount of differentially methylated cytosines within the primer regions. Based on two nanopore sequencing data sets we compared the results of `diffONT` to `metilene`, `DSS` and `pycoMeth`. We show that the regions predicted by `diffONT` are more specific towards hypermethylated regions and more usable for MSP. `diffONT` accelerates the design of methylation-specific diagnostic assays, bridging the gap between theoretical research and clinical application.

## Introduction

For the detection of genome-wide DNA methylations different methods have been used over the last decades, including bisulfite sequencing, DNA digestion with methylation-sensitive restriction enzymes, and affinity purification of methylated DNA combined with array hybridization (1). Each method has its specific applications, advantages, and limitations (1). Bisulfite treatment (2), the gold standard for detecting methylated cytosines (3), offers high-resolution mapping of methylation sites but can be labor-intensive and costly. Furthermore, bisulfite treatment is harsh and requires a high amount of input DNA. Digestion with methylation-sensitive restriction enzymes provides a more straightforward approach but lacks the resolution and comprehensiveness of bisulfite sequencing. Affinity purification of methylated DNA, on the other hand, enables the enrichment of methylated regions but may introduce biases depending on the antibody or binding protein used (4). Besides, it only indirectly resolves methylation patterns. The choice of method often depends on the specific research question, sample availability, and required resolution.

In cancer research, the accurate detection of DNA methylation is crucial for understanding tumorigenesis and for developing diagnostic and therapeutic strategies. One approach used in clinical diagnostic tests is methylation-specific PCR (MSP) (5), which uses bisulfite treatment, converting unmethylated cytosines to uracils while leaving methylated cytosines unchanged, allowing for the differentiation between methylated and unmethylated DNA by specific primers only binding to methylated sites. MSP is both simple and economical, making it suitable for population-wide screening (4). The design of MSP primers has to balance (a) specificity, to avoid creating unspecific PCR products or false positives in healthy individuals, and (b) efficiency, to achieve the theoretical maximum of duplication in each amplification step. Generally, a primer length of 18 to 24 nt is used, influencing specificity, with bisulfite-conversion-based PCR typically requiring longer primers (20 to 30 nt) due to lower sequence complexity(6; 7). PCR products of 150 to 1000 bp are produced during amplification, with shorter products being more specific; clinical assays advise 120 to 300 bp, and PCR products based on bisulfite treated DNA are advised to be below 400 nt(6; 7). The GC content of the PCR primers should be moderate, with 50-60% being ideal (8). For bisulfite-conversion-based PCR the primer sequence should also have enough non-CpG cytosines (5). Fig. 1 shows an example of an MSP primer pair design for a given DNA sequence.

**Fig. 1.**
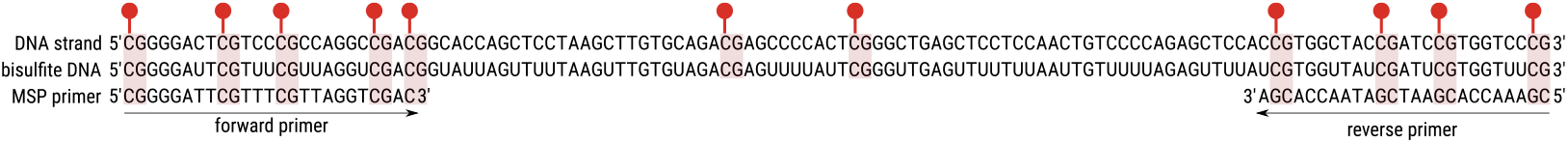
Standard primer design workflow, assuming all CpG dinucleotides are completely methylated. By treating the DNA with sodium-bisulfite (bisulfite), all non-methylated cytosines are deaminated into uracils. In a following PCR, the methylation specific primers bind only to methylated DNA stretches. Thus, only methylated DNA stretches are amplified, unmethylated DNA is not amplified. This allows a methylation-specific amplification of DNA. Primer length: 25 nt, PCR product length: 120 nt.

For the detection of marker regions for MSP, ONT sequencing is advantageous as it considers the entire genome, providing a comprehensive, unprejudiced view beyond just the regions described in literature or known disease-associated genes. This method surpasses the limitations of focusing solely on known regions, offering a broader and more inclusive analysis, while also directly measuring DNA methylation without the need of an amplification or conversion step on a single nucleotide resolution (9). While nanopore sequencing is not applicable as a diagnostics means in most applications due to its lack of economy and sensitivity, it is well-suited for detecting differentially methylated regions using tools developed for short or long-read sequencing (10; 11; 12; 13). These regions could potentially serve as targets for MSP.

A variety of bioinformatic tools are available for measuring DNA methylation differences, primarily designed for short reads or bisulfite data analysis:

`DSS` (Dispersion Shrinkage for Sequencing data) (14) is specifically designed to accommodate a limited number of biological replicates. It has an integrated beta-binomial regression model with an ’arcsine’ link function, utilizing a Bayesian hierarchical framework for differential methylation analysis. It assesses per-cytosine significance using a Wald test. However, `DSS` has been noted in a comparative study for its slower computational speed and potential memory issues, as well as a tendency to identify regions that may exceed the actual size of DMRs (15). It has also been observed to have lower precision when dealing with small number of replicates, with a notable drop in precision observed for n=3 compared to n=10 (10; 16).

`metilene` (11) is designed for high-speed performance and efficient memory usage (17; 11) to efficiently process large datasets. Additionally, it has been reported to work well with low coverage data. It has a binary segmentation algorithm implemented, coupled with a two-dimensional statistical test for detecting DMRs. While `metilene` is not effective for analyzing methylation differences in single-CpG analysis (16), it is known to identify DMR boundaries more accurately than `DSS` (16). However, it has been noted to identify numerous DMRs with low mean methylation differences between groups, although this parameter can be adjusted (15). Additionally, when applied to nanopore sequencing data, the data needs to be formatted to be in an adequate input format for `metilene`.

`pycoMeth` (12) is to our knowledge the only tool designed for differential methylation analysis using nanopore sequencing data processed through `Nanopolish`. `pycoMeth` shows increased recall and precision in DMR testing, especially for detecting low effect-size methylation changes in low-coverage settings (12). Additionally, by utilizing Bayesian changepoint detection, `pycoMeth` accommodates methylation call uncertainties instead of binarization of methylation probabilities as the other tools described above (12). `pycoMeth` is suitable for comparing two or more samples. `pycoMeth` focuses on identifying methylated regions based on nanopore sequencing data. However, it fails to support newer methylation callers such as implemented in the current state-of-the-art basecaller `Dorado`^1^.

The various analysis tools for identifying differential methylation patterns differ significantly in performance, including speed, memory usage, number of sample groups to compare, coverage, and the ability to detect small effect sizes. However, most of these tools are not suitable for the analysis of nanopore data.

To date, no tool has been specifically designed for the adequate detection of primer binding sites or the prediction of potential biomarkers that can be easily and affordably detected using methylation-specific PCR (MSP). Especially, identifying DNA methylation marker regions that can be effectively translated into clinical practice remains a significant challenge (4). Biomarkers not only need to be clearly differentially methylated in disease but also must be suitable for clinical diagnostic tests, ensuring that healthy individuals are minimally misclassified as having cancer, by minimizing the number of false positive test results and at the same time maximizing detection rates of diseased individuals.

Here, we present the workflow `diffONT`, written in Python, which addresses critical gaps in DNA methylation marker discovery and validation, particularly for clinical diagnostic via MSP. `diffONT` predicts potential diagnostic biomarkers based on low-coverage nanopore sequenced whole genome DNA methylation data independent of the used methylation caller, considering clinical constraints by minimizing false positives in the prediction of candidate regions and applying it to clinical datasets. `diffONT` allows for customization of parameters such as methylation thresholds, region length, minimum differential CpG sites, and PCR product size. Using an Oxford Nanopore Technology (ONT) dataset for validation, our tool has been benchmarked against established methods like `metilene`, `DSS`, and `pycoMeth`, demonstrating fewer regions detected, however, with more restrict methylation differences and direct applicability for MSP. This approach not only explores upstream methylation patterns but also considers methylation dynamics across diverse genomic contexts, thereby enhancing the quality and relevance of identified biomarkers for clinical applications.

## Results

In this study, we demonstrate the functionality of `diffONT`, a tool designed to predict methylation-specific PCR biomarkers from Nanopore-sequenced DNA data, using a publicly available ONT dataset. We describe the identification of CpGs of interest, followed by the definition of potential primers and the resulting MSP regions. To validate the effectiveness of our approach, we benchmark `diffONT` against other tools with similar functionality, namely `metilene`, `DSS`, and `pycoMeth`.

### Non-uniform distribution of CpGs of interest

In order to identify biomarker candidates for clinical diagnostics, we first determined the CpG sites of interest according to Fig. 9A-G. These have to be nearly unmethylated in control samples (less than 10 % methylation) and show significant hypermethylation in disease samples.

For the ONT dataset analyzed, about 3 874 242 CpGs (13,4 % of CpGs in the reference genome) of the complete human genome are predicted to be differentially methylated, see Fig. 2A. The distribution of CpGs of interest among the chromosomes generally correlates with chromosome length (SFig. S1B), with more CpGs of interest on the longer chromosomes 1 to 5 and fewer on the shorter ones, 19 to 22. Surprisingly, chromosome X forms an exception: despite being similar in size to chromosome 7, it contains 31.7 % more CpGs of interest than expected. To understand the unexpected behavior of chromosome X, we calculated for each chromosome the absolute number of (i) genes, (ii) CpGs, and (iii) CpG islands (SFig. S2A-C), as well as these attributes normalized by chromosome length (SFig. S2D-F). While these factors do not explain the large number of CpGs of interest, it was notable that chromosome X had comparatively low coverage, see STab. S2, which might suggest an artifact. A possible explanation might be abnormalities in the chromosomal copy numbers in the male cell line COLO829 (21), including, a duplication of the X chromosome. To determine whether the CpGs of interest are clustered on the chromosome or evenly distributed, we applied a sliding window of 10 000 nt across the entire genome (for comparison, the average CpG island is 700 nt long). Our analysis revealed that most windows have a ratio of CpGs of interest to total CpGs below 25 %, with only a few windows showing a ratio above 70 % CpGs of interest, see Fig. 2B. A chromosomal distribution can be found in SFig. S3A and choosing different window sizes does not change the observed effect, see SFig. S3B. A scatter plot reveals several interesting windows with a high percentage of CpGs of interest and a substantial total number of CpGs within the windows, see Fig. 2C. Notably, these windows overlap with the genes HIC1, GATA2, and TSPOAP1. Among these, HIC1 stands out as a tumor suppressor gene and features a CpG island spanning its promoter region (18). Also, GATA2 and TSPOAP1 show notable associations with cancer: GATA2 is implicated in hematological malignancies and potentially influences tumor microenvironment interactions (19), while TSPOAP1, involved in mitochondrial function and stress responses may affect tumor cell metabolism and behavior, highlighting their relevance in the dataset comprising normal B lymphoblasts (COLO829BL) and melanoma fibroblasts (COLO829) (22; 20). The fact that a known tumor suppressor gene stands out among the analyzed windows, in relation to a high percentage of CpGs of interest highlights the potential of our approach to identify relevant biomarkers in cancer.

**Fig. 2.**
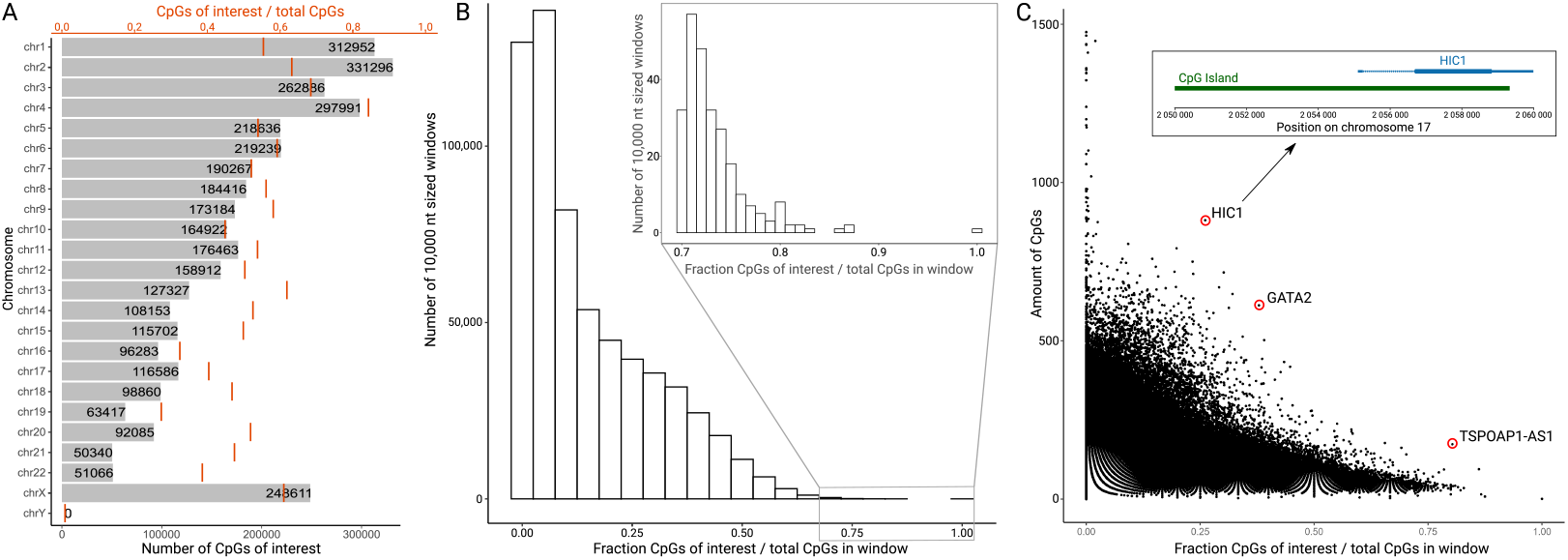
(A) Distribution of CpGs of interest across human chromosomes. As expected, the distribution largely correlates with chromosome length; however, chromosome X contains more CpGs of interest than anticipated. The relative amount of CpGs of interest compared to the total number of CpGs per chromosome vary less, with low fractions on chromosomes 17 and 19 and a high fraction of CpGs of interest on chromosome 4. (B) Histogram showing the distribution of the ratio of CpGs of interest to total CpGs across the human genome. Ratios were calculated using 10 000 nt windows, shifted by half the window length, and binned into 0.05 increments. For ratios above 0.7, data was rebinned into 0.01 increments. The highest ratio (1.0) is on chromosome 9, with only one CpG of interest. The second highest (0.88) is on chromosome X, with seven CpGs of interest out of eight. (C) Scatter plot depicting the ratio of CpGs of interest to total CpGs for various windows, with the total number of CpGs indicated on the vertical axis. Outlier points with a high percentage of CpGs of interest and a high amount of CpGs for the window are marked and overlap the genes HIC1, GATA2, and TSPOAP1, which are known to be associated with cancer (18; 19; 20).

CpG islands are regions of the genome rich in CpG dinucleotides and are often associated with gene promoters and regulatory regions (23). About 50 % of CpG islands contain known transcription start sites (24), particularly most housekeeping genes have CpG island covering their transcription start (25; 26). In our analysis, we found that, on average, only 5 % of the CpGs of interest overlap with these CpG islands, see Tab. 1. Independent analysis of each chromosome reveals considerable variability, with overlaps ranging from 2.04 % on chromosome 4 to 16.79 % on chromosome 19. This variability does not align with the overall distribution of CpG islands across chromosomes, see SFig. S2C. Notably, some chromosomes exhibit a higher overlap than others, suggesting that specific regions might influence the presence of CpGs of interest.

**Table 1.** Overview of the percentage overlap between predicted CpGs of interest, primers, and MSP regions with CpG islands. The overlap increases with each pipeline step, from an average 5 % for CpGs to an average of 72 % for MSP regions.

| Chr | CpG | Primer | MSP | Chr | CpG | Primer | MSP |
| --- | --- | --- | --- | --- | --- | --- | --- |
| Overlap with CpG islands [%] |  |  |  | Overlap with CpG islands [%] |  |  |  |
| 1 | 4.72 | 51.02 | 63.98 | 13 | 2.68 | 43.99 | 85.00 |
| 2 | 3.52 | 45.20 | 83.20 | 14 | 4.62 | 40.49 | 39.00 |
| 3 | 3.51 | 49.88 | 87.69 | 15 | 4.66 | 41.22 | 41.80 |
| 4 | 2.04 | 41.21 | 43.84 | 16 | 6.70 | 47.44 | 80.00 |
| 5 | 2.87 | 48.18 | 78.24 | 17 | 10.68 | 57.89 | 84.84 |
| 6 | 3.70 | 40.23 | 62.15 | 18 | 2.50 | 42.86 | 92.31 |
| 7 | 5.05 | 52.64 | 78.74 | 19 | 16.79 | 58.64 | 72.77 |
| 8 | 3.32 | 45.04 | 87.80 | 20 | 6.27 | 51.02 | 53.70 |
| 9 | 4.66 | 45.87 | 66.44 | 21 | 4.41 | 42.96 | 93.06 |
| 10 | 3.79 | 38.86 | 81.06 | 22 | 10.62 | 61.38 | 97.75 |
| 11 | 4.59 | 49.69 | 79.29 | X | 2.84 | 50.14 | 94.15 |
| 12 | 4.19 | 46.67 | 76.26 | Y | 0.00 | 0.00 | 0.00 |

### Methylation-specific PCR primer designed by diffONT

For methylation-specific PCR, primers need to be 18-24 nt long, contain at least three CpGs of interest each, and receive a primer score reflecting the methylation difference between patient and control samples, see Fig. 9H/I. We provide a comprehensive list of potential primers, detailing their genomic position, length, number of CpGs of interest, score, and GC content.

Applying `diffONT` to the ONT dataset resulted in 16 837 primer regions distributed across all human chromosomes. The majority of primers are predicted on chromosomes 1, X, 2, 17, and 6, with 1 425, 1 396, 1 261, 1 097, and 1 039 primers identified, respectively, see Fig. 3A. Most potential primers of `diffONT` (13 329) contain the required minimum of three CpGs of interest and only one primer contains eight CpGs of interest, see Fig. 3B. Tab. 2 provides a summary of the CpGs of interest compared to the total number of CpGs per primer. Overall, the number of CpGs of interest tends to increase as the total CpG count per primer rises. However, the majority of primers still contain only three CpGs of interest, regardless of the maximum number of CpGs. Over 96 % of 24 nt-wide windows (250 561 293 out of 258 326 583) do not contain any CpG of interest. While only about 5 % of CpGs of interest overlap with CpG islands, approximately 50 % of primers overlap with CpG islands, see Tab. 1. It varies between 38.86 % on chromosome 10 and 61.38 % on chromosome 22.

**Fig. 3.**
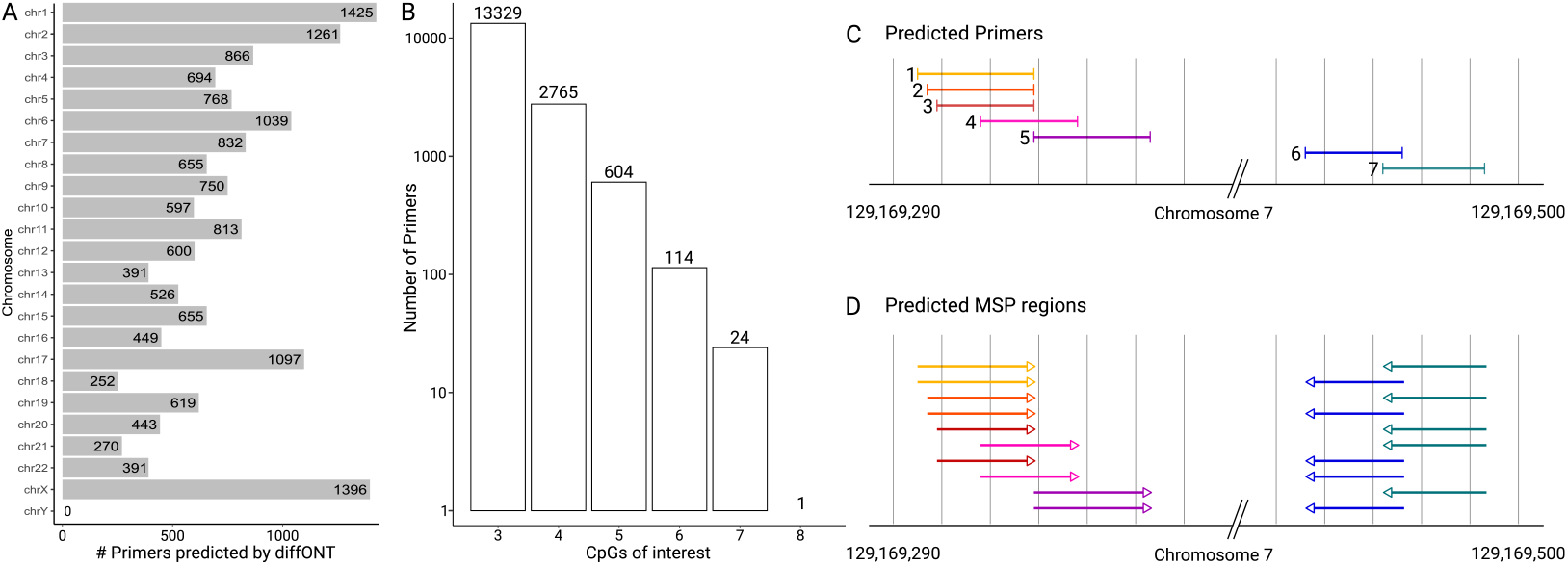
(A) Distribution of primers predicted by `diffONT` across chromosomes. The distribution generally follows chromosome length (SFig. S1 B), with exceptions on chromosomes 17 and X, which have more CpGs of interest than expected. (B) In total, 16,837 primers contain at least three CpGs of interest, with the majority of primers predicted by `diffONT` containing exactly three CpGs of interest. Note: the y-axis is visualized on a logarithmic scale (log10). (C) Most individual primers predicted by `diffONT` show overlap. (D) The various primer combinations generate multiple MSP regions with slightly different scores and characteristics, such as length and number of CpGs, as shown in Fig. 9K.

In CpG-rich regions, such as CpG islands, multiple overlapping primers are predicted (Fig. 3C). As multiple combinations of predicted primers are possible to yield a PCR product (within the range of 400 nt), multiple MSP regions containing the same primers are possible (Fig. 3D). The best MSP region is the combination of primers with the highest score (i.e. those primers with the highest coverage and non-control methylation).

**Table 2.** Distribution of primers (column 2) by the total number of CpGs (column 1) and the number of CpGs of interest (column 3). For example, among 1 342 primers containing 6 CpGs, 614 of those primers have 3 CpGs of interest, 434 have 4, 233 have 5, and 61 have all 6 CpGs of interest. Note that primers with a high number of total CpGs may be unsuitable due to elevated annealing temperatures.

| total<br>CpGs | Primer | Primers with $n$ CpGs of interest<br>($n$ from 3 to number of total CpGs) |
| --- | --- | --- |
| 3 | 7 831 | 7 831 |
| 4 | 4 336 | 3 092 - 1 244 |
| 5 | 2 808 | 1 599 - 921 - 288 |
| 6 | 1 342 | 614 - 434 - 233 - 61 |
| 7 | 432 | 164 - 144 - 67 - 40 - 17 |
| 8 | 78 | 25 - 22 - 15 - 12 - 3 - 1 |
| 9 | 9 | 4 - 0 - 1 - 1 - 3 - 0 - 0 |
| 10 | 0 | 0 - 0 - 0 - 0 - 0 - 0 - 0 - 0 |
| 11 | 1 | 0 - 0 - 0 - 0 - 1 - 0 - 0 - 0 - 0 |

### MSP regions by diffONT

The best combination of primers is selected based on the highest summed score of two primers with a user-defined distance (60-400 nt), see Fig. 9J/K. `diffONT` presents a list of MSP regions containing score, coverage, and length. In total, the `diffONT` algorithm reports 6 745 potential MSP regions for the ONT dataset across the human genome. The highest number of MSP regions were predicted on chromosome X with 1 146 predicted MSP regions (see Fig. 4A). The fewest hits are located on chromosomes 21, 18, and 13 with 72, 78, and 100 MSP regions, respectively. No suitable MSP region is predicted on chromosome Y, as the Y chromosome is lost in COLO829. The ONT dataset shows differences in the coverage between the chromosomes, see STab. S2, with e.g., very low coverage (0.27 – 0.4 X) on chromosome Y for the melanoma fibroblast samples.

**Fig. 4.**
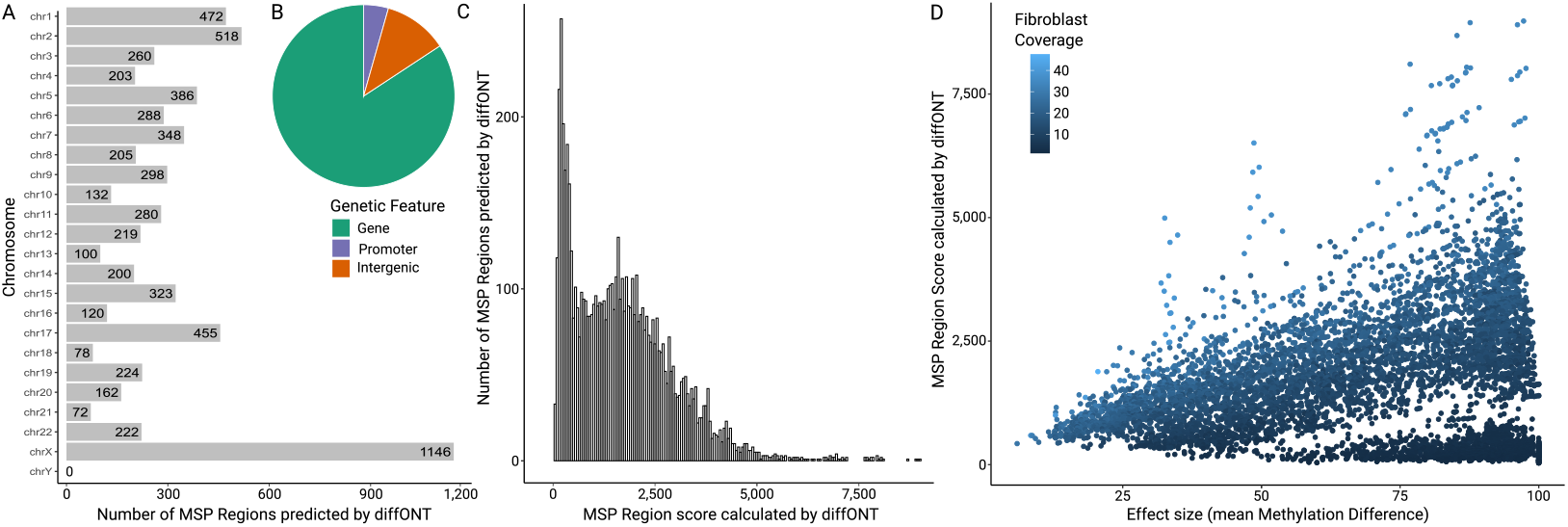
(A) Distribution of predicted MSP regions by `diffONT` on the ONT dataset across chromosomes. The regions are unevenly distributed, with the highest concentrations on chromosomes X, 1, 2, and 17. (B) MSP regions overlap with (annotated) genomic features: 5 683 genes, 296 promoter regions (-500 nt), and 766 intergeneic regions. (C) Histogram of `diffONT` score values for all MSP regions predicted on the ONT dataset, with scores binned in intervals of 50. Two distinct distributions are observed: one peaking around 200 and another around 2 000, with scores above 5 000 appearing as outliers. (D) Visualization of MSP region scores versus mean methylation differences between melanoma fibroblast and normal B lymphoblast samples for all MSP regions predicted by `diffONT` on the ONT dataset. Color represents average coverage in melanoma fibroblast samples. Regions with low scores despite high methylation differences (bottom right) exhibit low coverage in melanoma fibroblast samples.

However, the mean coverage of the predicted regions follows a similar bimodal distribution for both fibroblast and normal lymphoblast samples, see SFig. S1A.

Most of the 6 745 MSP regions overlap with annotated genomic features, as shown in Fig. 4B. Specifically, 5 683 MSP regions (84 %) overlap with an annotated gene, another 296 regions (4.39 %) overlap with the 500 nt upstream of a gene and about 10 % (766 MSPs) are intergenic.

The score for the 6 745 predicted MSP regions ranges from 16.44 to 8 978.00, see Fig. 4C. The score values follow a bimodal distribution, with a peak below 500, created by 1 504 MSP regions, and a second peak at around 2 000 created by 5 138 MSP regions with a score between 500 and 5 000. Only 101 MSP regions have a score above 5 000.

The mean methylation difference values (also called effect size), generated by subtracting the average melanoma fibroblast samples’ methylation from the average normal B lymphoblast samples’ methylation, are shown in SFig. S4A reveals a high methylation difference (above 75 %) for about 50 % of the regions (3 306 out of 6 745). This is an interesting result, as the `diffONT` algorithm does not require high methylation differences. In contrast, only 317 of the predicted MSP regions have a mean methylation difference below 25 %.

Comparing the MSP region score with the mean methylation difference for all MSP regions predicted by `diffONT` reveals a positive correlation, see Fig. 4D. Regions with a higher mean methylation difference tend to have higher scores, likely because the score is influenced by the average disease methylation, and regions with significant methylation differences typically have higher disease methylation, see SFig. S4B . However, some regions exhibit a high mean methylation difference but a low score, which can be attributed to low coverage in the melanoma fibroblast samples. Since the score incorporates both disease sample methylation and coverage, low coverage results in a lower score, explaining this discrepancy.

A histogram of the average disease and control samples coverage shows a bimodal distribution of coverage values, with one peak at around 2 X coverage, and another one at around 15 X coverage, see SFig. S1A. We analyzed the overlap of the MSP regions predicted by `diffONT` with CpG islands, see Tab. 1. On average, 72 % of MSP regions overlap at least one CpG island by at least one nucleotide. However, the overlap varies significantly across chromosomes: only 39 % and 42 % of MSP regions on chromosomes 14 and 15, respectively, overlap with CpG islands, while over 90 % of MSP regions overlap CpG islands on chromosomes 18, 21, 22, and X. Interestingly, while the average overlap with CpG islands across all chromosomes is about 5 % for individual CpGs of interest, it increases to 46 % for primer regions and to 72 % on MSP regions.

The predicted region with the highest score overlaps an annotated CpG island as well as the 3’ UTR of the gene TSPAN33 (Tetraspanin 33), see Fig. 5, a gene for which differential methylation is linked to several lymphomas (27). The methylation for the predicted region in the TSPAN33 3’ UTR strongly differs between the melanoma fibroblast and normal B lymphoblast replicates.

**Fig. 5.**
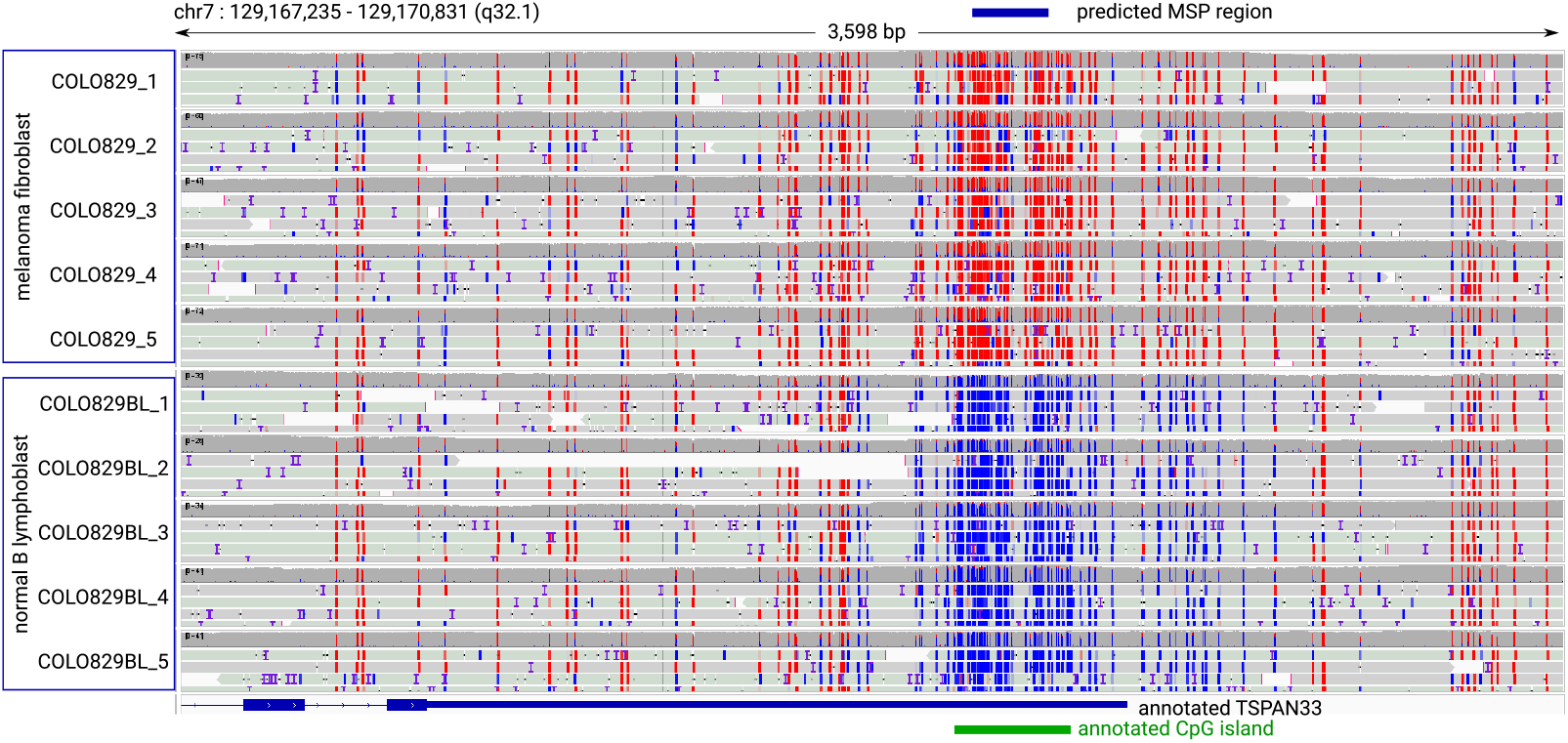
Visualization of the highest-scoring MSP region predicted by `diffONT` on the ONT dataset using the Integrative Genomics Viewer (IGV). The predicted `diffONT` region is annotated in blue at the top, with the gene TSPAN33 and the associated CpG island annotated below (blue and green, respectively). The 5mC methylation status is color-coded: red for methylated and blue for unmethylated sites. The melanoma fibroblast sample (5x) is displayed above the normal B lymphoblast sample (5x), showing strong methylation differences overlapping the TSPAN33 3’ UTR.

Another region, which is among the 50 highest-scored MSP regions overlaps the 5’ UTR of transcription factor TCF20, see Fig. S4C and a predicted CpG island. Variants of TCF20 are known to impair neurodevelopmental disability (28). Additionally, TCF20 is known to be abundantly expressed in small cell lung cancer (29) and to distinguish desmoid tumors from nodular fasciitis (30). The predicted MSP region shows a higher amount of CpG dinucleotides than the surrounding genomic region and a high methylation difference between the fibroblast and lymphoblast samples. The top 10 ranked MSP regions can be found in STab. S3.

#### Performance benchmark of diffONT

To the best of our knowledge, none of the existing tools predicts MSP regions for PCRs in clinical applications. Instead, all currently available tools screen for differentially methylated regions (DMRs) in general. That is why we compared the MSP regions of our tool `diffONT` with DMRs predicted by `metilene`, `DSS`, and `pycoMeth`. We expect our MSP regions to form a subset of DMR regions, due to our stringent selections based on (i) an applicability in a clinical context, i.e., a high methylation in disease samples and almost no methylation in control samples, instead of finding general differences in methylation; and (ii) a high methylation rate within only 24 nt allowing primer design for methylation-specific PCR.

Clinical nanopore sequencing data often has limited coverage (∼7 X per replicate per flow cell for the dataset used). We focus our benchmarking on `metilene`, which performs well with such data (11), and `DSS`, which is currently recommended by ONT for the analysis of nanopore sequencing data. Additionally, we show benchmarking data for `pycoMeth`, as this tool has been developed especially for DMR detection in nanopore sequencing data.

### Comparison to metilene

`metilene` was implemented to study DNA methylation differences, e.g., after environmental stress in plants (31), during development (32; 33), and for the prediction of cancerous lung tissue (34). We applied `metilene` to predict DMRs to the same ONT dataset analyzed with `diffONT`. `metilene` predicted 621 523 DMRs, of which only 2 342 overlap with MSP regions of `diffONT`, see Fig. 6A. Contrarily, 6 032 (98 %) of 6 745 MSP regions predicted by `diffONT` overlap with a DMR predicted by `metilene`. There are mainly two differences between the regions predicted by `diffONT` and `metilene`: (i) the length of `metilene`’s DMRs ranges from 10 nt to 7 486 nt, whereas `diffONT`’s MSP regions range from 60 nt to 400 nt, see Fig. 6B, resulting in generally longer regions predicted by `metilene`, additionally, (ii) `diffONT` only considers up-regulated methylation patterns compared to control samples, whereas `metilene` predicts hyper- and hypomethylated regions (Fig. 6C, green). These two differences explain the low fraction of DMRs not intersecting at all with MSP regions.

**Fig. 6.**
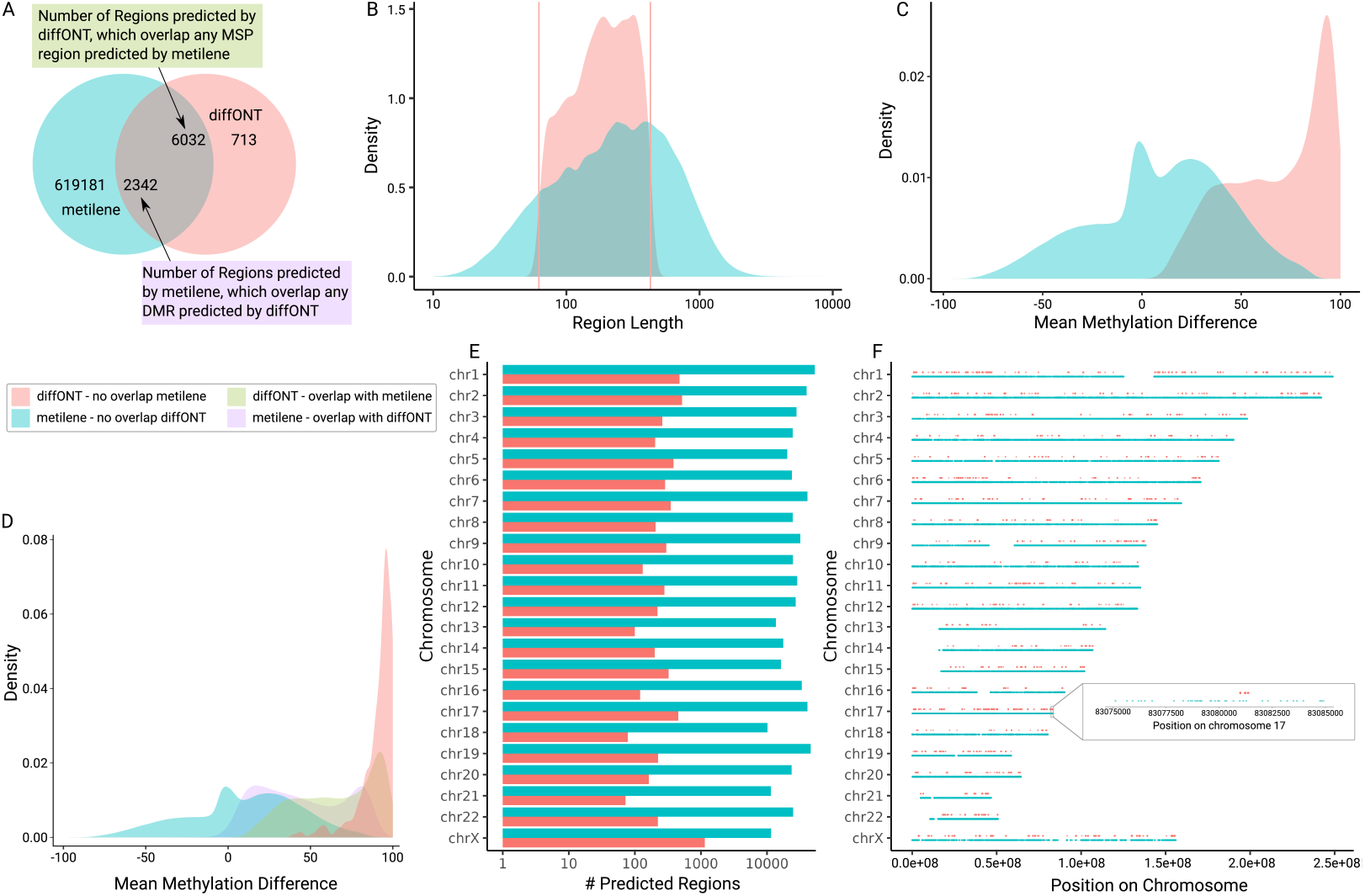
Comparison between `diffONT` (red) and `metilene` (green) for the ONT dataset, comparing cancerous fibroblast cells with normal lymphoblast cells. (A) Venn diagram showing the overlap of predicted regions. (B) Frequency density of the region length. All regions returned by `diffONT` range from 60 to 400 nt (vertical lines) due to the given values for minimum and maximum amplicon length in the algorithm, see Fig. 9J. Please note: x-axis is shown in log(10) scale. (C) Frequency density of the mean methylation differences between the melanoma fibroblast and normal lymphoblast replicates. (D) The distribution of the four fields of the Venn diagram. (E) Number and (F) distribution of DMR (from `metilene`) and MSP (from `diffONT`) regions per chromosomes.

The density distribution of mean methylation differences of the regions predicted by `metilene` ranges from -100 to 100. Interestingly, a peak around -1 may highlight regions relevant for studying subtle methylation changes, as needed e.g. in early disease stages. This is, however, not applicable for methylation-specific PCRs in a clinical context. The regions predicted by `diffONT` all have a methylation difference peak at around 93, thus `diffONT` focuses on regions with high methylation differences. Notably, for all the comparisons made, the `diffONT` score filter has not been applied at this stage, thus, applying it will further increase the rate of methylation differences among the finally selected MSP regions, leading to even more accurate results of `diffONT`. In order to understand the distribution of methylation differences in more detail, we further distinguished the four fields of the Venn Diagram from Fig. 6A in Fig. 6D. Due to the specific design of `diffONT` for clinical applicability, which limits the analysis to regions where control samples exhibit little to no methylation, most `diffONT` results display a very high mean methylation difference. Accordingly, the `diffONT` results can be better applied in the clinical context.

The distribution, number, and appearance of `diffONT` and `metilene` regions across the chromosomes are reported in Tab. 3, Fig. 6E (note the log-scaled x-axis) and Fig. 6F. Both tools predict some chromosomal parts to have no methylation differences, such as the 5’ end of acrocentric chromosomes 13, 14, 15, 21, and 22.

**Table 3.**
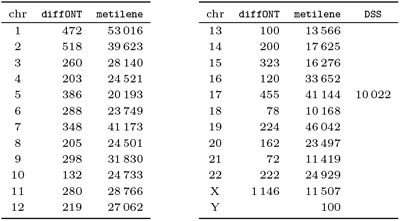
Number of regions called per chromosome by diffONT and metilene on the ONT dataset. For both methods the distribution of predicted regions among the chromosomes is quite similar. Both methods predict most regions on chromosomes 1 and 2 and the least regions on chromosome 21. Low counts also exist on alternative chromosomes (data not shown).

| chr | diffONT | metilene | chr | diffONT | metilene | DSS |
| --- | --- | --- | --- | --- | --- | --- |
| 1 | 472 | 53 016 | 13 | 100 | 13 566 |  |
| 2 | 518 | 39 623 | 14 | 200 | 17 625 |  |
| 3 | 260 | 28 140 | 15 | 323 | 16 276 |  |
| 4 | 203 | 24 521 | 16 | 120 | 33 652 |  |
| 5 | 386 | 20 193 | 17 | 455 | 41 144 | 10 022 |
| 6 | 288 | 23 749 | 18 | 78 | 10 168 |  |
| 7 | 348 | 41 173 | 19 | 224 | 46 042 |  |
| 8 | 205 | 24 501 | 20 | 162 | 23 497 |  |
| 9 | 298 | 31 830 | 21 | 72 | 11 419 |  |
| 10 | 132 | 24 733 | 22 | 222 | 24 929 |  |
| 11 | 280 | 28 766 | X | 1 146 | 11 507 |  |
| 12 | 219 | 27 062 | Y |  | 100 |  |

In terms of run time `diffONT`, though with a shorter pre-processing time, performs much slower than `metilene`. However, `diffONT` can predict highly methylated regions more precisely, see Tab. 4. `diffONT` is computationally even more intensive for whole-genome analysis and more efficient for smaller regions, such as individual chromosomes. The longer runtime is initially counterintuitive given the smaller number of regions `diffONT` predicts. It arises, however, due to single nucleotide analysis and calculation of single nucleotide statistics (Fig. 9E).

**Table 4.**
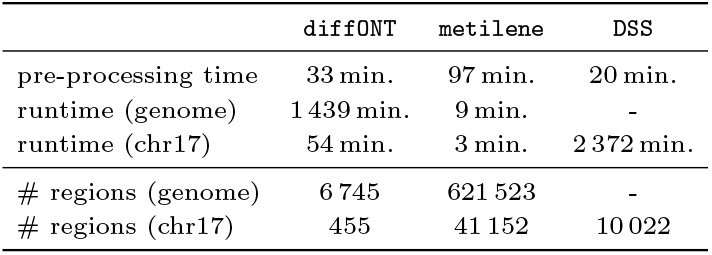
Comparison of the main result statistics between diffONT, metilene, and DSS on the ONT benchmarking dataset. Time calculated for pre-processing starting with (unsorted) bedmethyl files. # regions – number of DMRs and MSP regions, respectively.

|  | diffONT | metilene | DSS |
| --- | --- | --- | --- |
| pre-processing time | 33 min. | 97 min. | 20 min. |
| runtime (genome) | 1 439 min. | 9 min. | - |
| runtime (chr17) | 54 min. | 3 min. | 2 372 min. |
| # regions (genome) | 6 745 | 621 523 | - |
| # regions (chr17) | 455 | 41 152 | 10 022 |

However, we can assume `diffONT` to predict hypermethylated regions more precisely than `metilene`, with higher accuracy.

### Comparison to DSS

`DSS` is widely used in various publications for detecting differentially methylated regions (DMRs) based on nanopore sequencing data (35; 13; 36) and is also recommended by ONT. However, for whole genome data analysis, `DSS` is notably slow, see Tab. 4. Therefore, in this work, `DSS` was applied to chromosome 17 only, which contains many regions predicted by both `diffONT` and `metilene`, see Fig. 6. The runtime of `DSS` for only one chromosome exceeds that of `diffONT` by factor *>* 40. For the whole genome, we allowed `DSS` to run for over three months before halting the process without any result.

`DSS` results of chromosome 17 could be filtered by p-value or mean methylation difference. The complete (unfiltered) number of reported DMRs was used for comparison. In total, `DSS` predicted 10 022 DMRs for the ONT dataset on chromosome 17, which is 20 times the number of regions predicted by `diffONT` (455) and about a quarter of the DMRs predicted by `metilene` (41 144), see Tab. 3.

The majority of `diffONT` regions (345 of 455, 76 %, Fig. 7A) were found by both tools, another 107 (24 %) by `metilene` but not `DSS`, and only 2 regions by `DSS` but not `metilene`. Similar to `metilene`, also `DSS` detects hyper- and hypomethylated regions, which results in a high number of regions (ca. 7 500) found by both tools but not `diffONT`. This is inherent to `diffONT`, being designed to detect hypermethylated regions only, which additionally requires almost no methylation in control samples for the following application of MSP. The coverage of the chromosome for DMR and MSP regions is for `diffONT` 0.04 %, for `metilene` 14.45 %, and for `DSS` 2.28 %, see Fig. 7B. The DMRs predicted by `DSS` have a required minimum length of 51 nt, and no maximum length, resulting in DMRs ranging from 51 nt to 3 331 nt with a median region length of 122 nt. Nevertheless, `DSS` yields a high number (2 150) of long DMRs exceeding 250 nt. The region length distribution of `DSS` is similar to the one for regions predicted by `metilene`, with DMRs in the range of 10 nt to 3 576 nt and a median region length of 209 nt, Fig. 7C. Regions predicted by `diffONT` on the other hand have a minimum and maximum length defined (60 nt and 400 nt, respectively) and have a larger median region length of 182 nt. Expecting hypermethylation in the disease samples, `diffONT` screens exclusively for MSP regions with a positive mean methylation difference, Fig. 7D. In contrast, both `DSS` and `metilene` screen for DMRs with both positive and negative mean methylation differences. Similar to the entire human genome, `metilene` predicts numerous DMRs with a mean methylation difference near 0, whereas `DSS` predicts very few DMRs with small methylation differences. It can be summarized, that both `DSS` and `metilene` predict much more and longer regions on chromosome 17, compared to `diffONT`, however, for the application of MSP in the clinical context, `diffONT` seems more suitable.

**Fig. 7.**
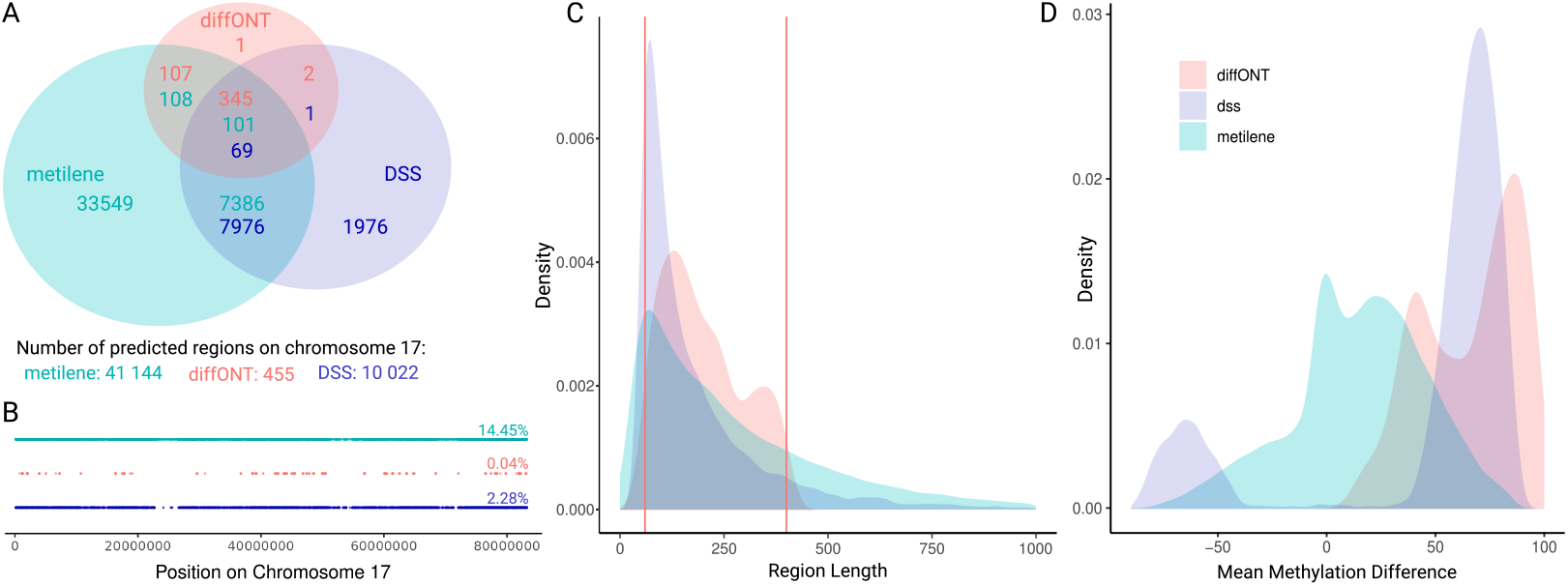
Comparison of `diffONT`, `metilene`, and `DSS` on chromosome 17 using the ONT dataset. (A) Venn diagram of predicted regions found by `diffONT`, `metilene`, and `DSS`. The intersections contain numbers in different colors which refer to the different tools. For example, 7 976 `DSS` regions overlap with `metilene` regions, whereas only 7 386 `metilene` regions overlap with `DSS` regions. (B) Distribution of regions predicted by `diffONT`, `metilene`, and `DSS` among chromosome 17 and coverage of chromosome by the DMRs and MSP regions predicted by `diffONT`, `metilene`, and `DSS`. (C) Density distribution of the length of predicted regions by `diffONT`, `metilene`, and `DSS`. All regions returned by `diffONT` range from 60 to 400 nt (vertical lines) due to the given values for minimum and maximum amplicon length in the algorithm (see Fig. 9J). Data is shown only for region length below 1 000 nt. (D) Density distribution of the mean methylation differences between the melanoma fibroblast and normal lymphoblast replicates for `diffONT`, `metilene`, and `DSS`.

#### Comparison with DMR calling tool pycoMeth

`pycoMeth` was the first tool being developed to predict DMRs from nanopore data, which includes storage, management and analysis of ONT DNA methylation data (12). Additionally, to our knowledge, `pycoMeth` is currently the only tool using low-coverage (up to 15 X) long-read nanopore sequencing data for predicting DMRs. For the comparison of `diffONT` to `pycoMeth`, both tools were applied to a male and a female sample, resulting in more than 100 000 predicted regions for each of the tools, Tab. 5.

**Table 5.** Distribution of regions predicted by diffONT and pycoMeth on the GIAB dataset. In total, 104 805 and 157 660 regions were identified for `diffONT` and `pycoMeth`, respectively. No regions are predicted on chromosome Y, since female cells were compared to male cells. Low counts also exist on alternative chromosomes (data not shown).

| chr | diffONT | pycoMeth | chr | diffONT | pycoMeth |
| --- | --- | --- | --- | --- | --- |
| chr1 | 1 235 | 11 117 | chr13 | 239 | 5 927 |
| chr2 | 530 | 12 153 | chr14 | 196 | 5 813 |
| chr3 | 509 | 9 662 | chr15 | 300 | 4 522 |
| chr4 | 331 | 8 944 | chr16 | 873 | 5 036 |
| chr5 | 577 | 9 383 | chr17 | 483 | 5 860 |
| chr6 | 435 | 8 783 | chr18 | 202 | 4 224 |
| chr7 | 199 | 8 814 | chr19 | 375 | 5 196 |
| chr8 | 571 | 7 441 | chr20 | 256 | 3 372 |
| chr9 | 251 | 7 523 | chr21 | 575 | 2 520 |
| chr10 | 359 | 8 641 | chr22 | 202 | 2 373 |
| chr11 | 266 | 8 190 | chrX | 95 527 | 4 346 |
| chr12 | 282 | 7 797 | chrY | — | — |

`pycoMeth` segments the genome based on methylation information into intervals, resulting in 584 171 intervals for the GIAB dataset. Differential methylation analysis of these intervals identified 157,660 intervals as DMRs, characterized by significant p-values. While about half of the MSP regions (54 122 out of 104 805) predicted by `diffONT` overlap a region predicted by `pycoMeth`, only a small fraction of regions predicted by `pycoMeth` (422 out of 157 660) are overlapping an MSP region, Fig. 8A. At first glance, this may appear to be a disproportionate distribution; however, it can be explained by the following observation: While the MSP regions predicted by `diffONT` have a defined length of 60-400 nt, the intervals predicted by `pycoMeth` range from 0 to 2 348 466 nt in length, see Fig. 8B, with a peak at 4 596 nt. Note, that `DSS` and `metilene` have not identified any such long DMR, see Fig. 6,

**Fig. 8.**
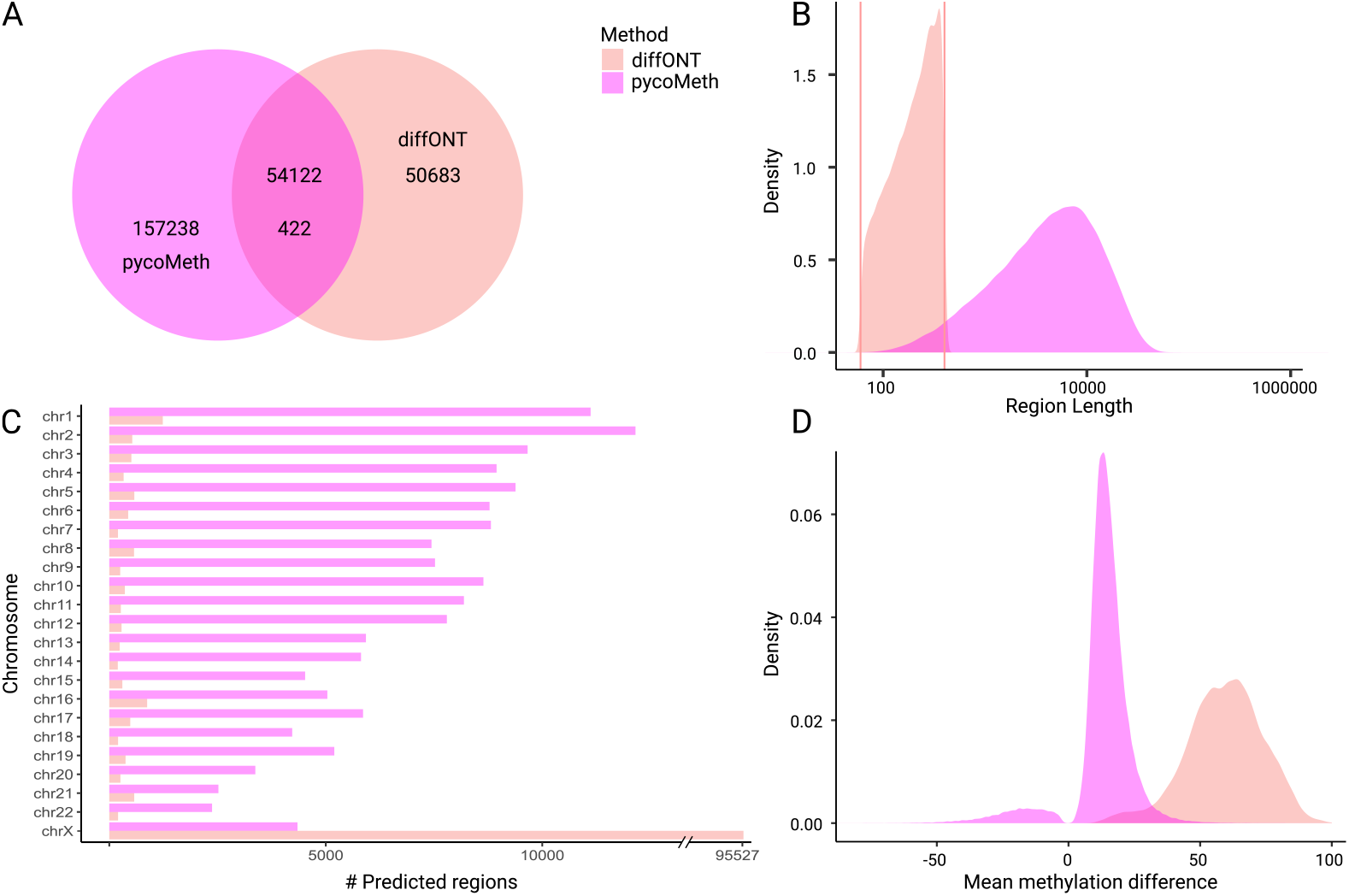
Comparison between `diffONT` (red) and `pycoMeth` (purple) for the GIAB dataset. (A) Venn diagram showing the overlap of predicted regions. (B) Density distribution of the region length of predicted regions by `diffONT` and `pycoMeth`. All regions returned by `diffONT` range from 60-400 nt (vertical lines) due to the given values for minimum and maximum amplicon length in the algorithm, see Fig. 9J. The X-axis is shown in log(10). The regions predicted by `pycoMeth` far exceed the length of those predicted by `diffONT`. (C) Distribution of the `diffONT` and `pycoMeth` predicted regions among the chromosomes. The X chromosome includes more than 95 500 (91 %) of the `diffONT` predicted regions. (D) Density distribution of the methylation difference of predicted regions by `diffONT` and `pycoMeth`. Methylation above 0 means higher methylation in the female sample HG004.

**Fig. 9.**
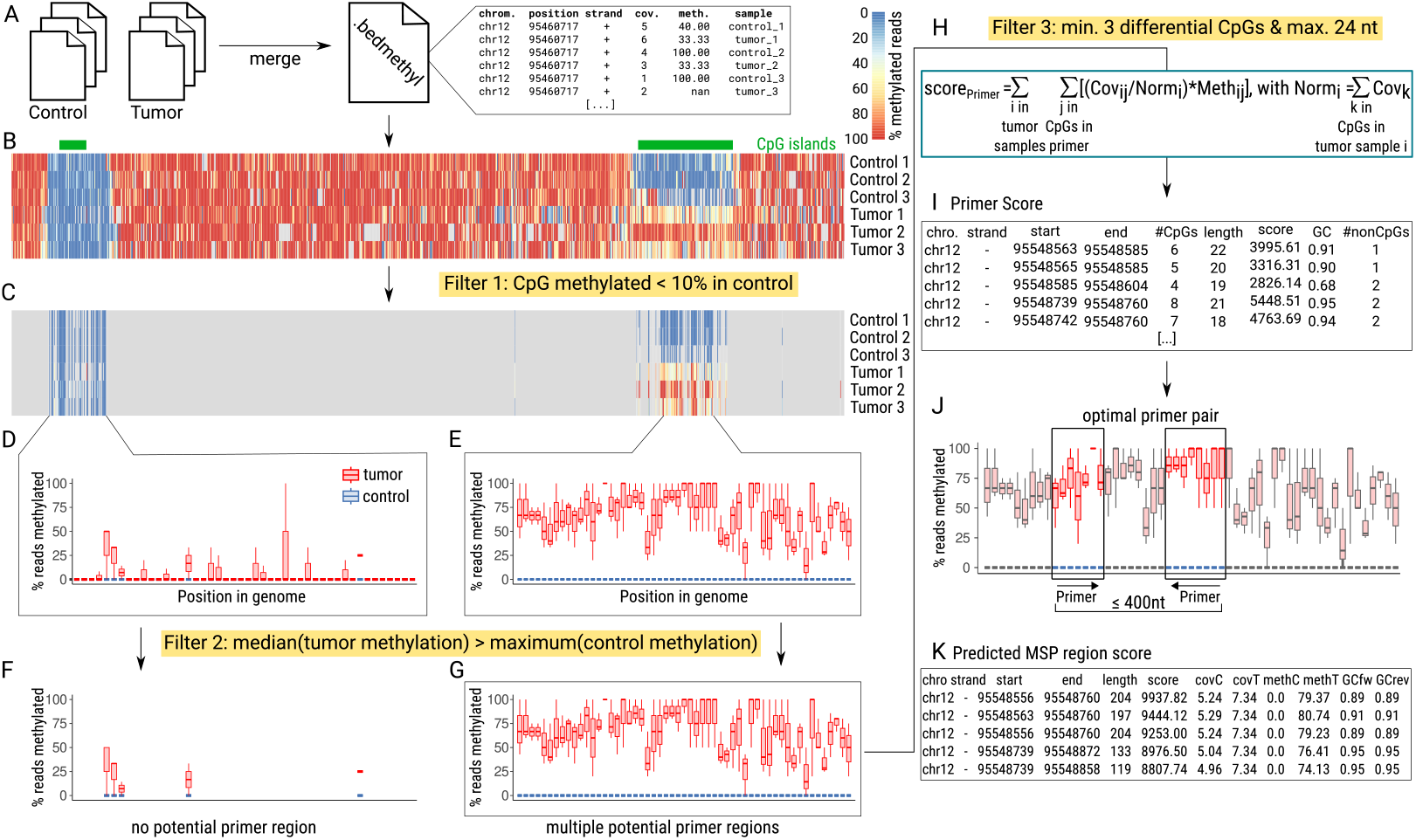
The `diffONT` workflow detects potential MSP regions, which differentiate between disease and control samples by containing CpG positions being unmethylated in control and methylated in disease samples (median methylation in disease samples above maximum methylation in control samples). Required input data is one `bedmethyl` file (e.g. generated by `modbam2bed`) per sample, containing information about number of reads and percentage methylation per cytosine for one sample. **(A)** In a preproccessing step `bedmethyl` files of different samples are merged and sorted by genomic position into one `bedmethyl` file. **(B)** Visualization of CpG methylation information for an exemplary region (exclusively (+) strand data is shown); `UCSC` CpG islands are annotated in green. **(C)** The methylation in the control samples is analyzed for each position. If the methylation exceeds the user-defined threshold (default *>*10 %) in only one of the control samples, the position is excluded from further analysis, Filter 1. **(D, E)** Boxplot statistics (0th, 25th, 50th, 75th and 100th percentile) are calculated for disease and control samples of each remaining position. These statistics are provided as tables to the user. **(F, G)** Only positions where the 50th percentile methylation in disease is above the 100th percentile methylation in control are considered as CpGs of interest. These are screened for potential primer regions. Primer regions should contain at least three differentiating CpGs within (per default) at most 24 nt. While for **(F)** the distance between the remaining CpGs is too large to form primers, for **(G)** multiple primers are possible. **(H)** For all potential primer regions a score is calculated, by summing up the normalized coverage multiplied with the percentage methylation for all remaining cytosine positions within the primer for all disease samples. **(I)** The table of potential primers and their scores is provided to the user. **(J)** The best combination of primers is selected based on the highest summed score of two primers within a user-defined region (default 60-400 nt). **(K)** The score for these MSP regions is determined by summing up the scores of the individual primers. The list containing predicted MSP region scores, GC content, coverage, and MSP region length is provided to the user.

Fig. 7. If many short `diffONT` MSP regions overlap with one long `pycoMeth` DMR region, then this results in a high overlap of MSP regions with DMRs and a small number of DMRs overlapping with any MSP result. The number of MSP regions predicted by `diffONT` for the GIAB dataset is approximately 15 times higher than the number predicted for the ONT dataset, generally indicating more methylation differences between males and females than between healthy and cancerous male cells. Interestingly, the MSP regions predicted by `diffONT` are distributed unequally among the chromosomes. An extremely high amount of predicted regions can be seen on chromosome X (95 527, ∼91 %), with the second-highest amount of MSP regions predicted on chromosome 1 (1 235), Fig. 8C. This can be explained by the fact, that DNA methylation on the X chromosome reflects sex-specific dosage compensation driven by X-chromosome inactivation (XCI) in the female sample, which has two X chromosomes (37).

Most of the DMRs predicted by `pycoMeth` are located on chromosome 2 (12 153) and chromosome 1 (11 117) (Fig. 8C, Tab. 5). `pycoMeth` does not predict an increased amount of DMRs on chromosome X (4 346). Instead, the distribution of DMRs predicted by `pycoMeth` follows the chromosome length distribution, SFig. S1. As `diffONT` is designed to show methylation differences detectable in an MSP, `diffONT` screens for methylation differences in one direction only, originally higher methylation in tumor samples compared to control samples. `pycoMeth` on the other hand, is not designed for this specific approach and thus predicts both hyper- and hypomethylated regions. As Fig. 8D displays, all `diffONT` predicted MSP regions are hypermethylated, i.e., showing higher methylation levels in the female sample. The majority of these predicted MSP regions display a methylation difference exceeding 50 %. As expected, `pycoMeth` predicts regions with both positive and negative methylation differences between the samples. However, most regions predicted by `pycoMeth` show a methylation difference below 50 % and thus are difficult to use in an MSP.

For comparison of runtime, we excluded basecalling and methylation calling. Running the `pycoMeth` workflow took 31 590 min. in total. The workflow consists of the short step `meth5 create m5`, followed by the methylation-based segmentation of the genome `pycometh Meth Seg`, which took 31 293 min., and is advised to be performed chromosome-wise. The final step `pycometh Meth Comp` took 104 min. Despite a longer preprocessing time (421 min.), `diffONT` is in total much faster than `pycoMeth` (1216 min.), see Tab. 6.

**Table 6.** Comparison of the main result statistics for diffONT and pycoMeth on the GIAB dataset.

|  | **diffONT** | **pycoMeth** |
| --- | --- | --- |
| pre-processing time | 421 min. | 0 |
| runtime | 795 min. | 31 590 min. |
| # predicted regions | 104 773 | 157 637 |

## Conclusion and Outlook

Here we presented `diffONT`, a tool to predict methylation-specific PCR biomarker regions, based on nanopore methylation sequencing data. `diffONT` is easily executable via the command line. In this study, we demonstrated that the MSP regions predicted by `diffONT` form a subset of the DMRs identified by other differential methylation calling tools, such as `DSS` and `metilene`. However, this subset has the advantage of showing only distinctly hypermethylated regions compared to the control sample, which is restricted to be nearly unmethylated. While `diffONT` is slower than `metilene`, its unique advantage lies in its ability to directly generate regions suitable for primer design in MSP. This feature makes `diffONT` particularly valuable for clinical applications, such as identifying biomarkers for cancer detection and potentially screening. Designed with a focus on this clinical use, `diffONT` simplifies and accelerates the development of methylation-specific diagnostic tests for cancer. By combining ease of use with direct applicability, `diffONT` provides a practical and efficient tool for bridging the gap between epigenomic research and clinical diagnostics.

Currently, `diffONT` does not incorporate allele-specific data (13) or information regarding the primer annealing temperature. However, this design choice allows users the flexibility to apply their own tools and methods for selecting primers, depending on their specific needs and preferences. Additionally, while `diffONT` is efficient in identifying MSP regions, its runtime could be further optimized by combining it with other DMR calling tools, such as `metilene`. Given that `metilene` includes nearly all regions predicted by `diffONT`, an effective strategy would be to use `metilene` for a pre-selection step followed by `diffONT` for analysis of the pre-selected regions. This combination could significantly reduce runtime while maintaining the accuracy and specificity of the MSP regions identified. These improvements would further strengthen `diffONT`’s utility in clinical and research applications.

Overall, `diffONT` presents a unique solution for the clinical use-case of predicting MSP-specific primer regions and thus can help to transfer knowledge of methylation-specific (cancer) markers into clinical diagnostics.

## Material and Methods

### The diffONT pipeline

`diffONT` is a Python-based tool available for download on GitHub^2^, designed for analyzing DNA methylation differences between a control group (e.g., healthy individuals) and a disease group (e.g., patients with the same cancer type). It evaluates methylation variations across the entire genome, taking into account sequencing coverage and specifically identifying regions where methylation is absent in the control group but elevated in the disease group. `diffONT` outputs and scores potential biomarker regions independent of their location relative to known genes, including upstream, downstream, intergenic, and gene body regions. Additionally, it suggests primer sequences suitable for methylation-specific PCR (MSP) to facilitate experimental validation of identified potential biomarkers.

#### Data preprocessing

A merged and sorted `bedmethyl` file, containing data from at least one control and one disease sample (see STab. S1) including an additional column for the sample name, serves as the input for `diffONT` using the `--bedmethyl` parameter. The required input `bedmethyl` file can be generated with the script `preprocess.py`, which merges multiple `bedmethyl` files into one `bedmethyl` file, and sorts it by genomic positions. An additional column defines the origin of the entry for each line. The `bedmethyl` file contains for each sample for each position and for each strand information about coverage and methylation percentage (Fig. 9A/B).

#### Filter 1: Non-methylated CpGs in control samples

For the usage in a clinical context, we aim to reduce the number of false positives and require the control samples to show almost no methylation and diseased samples being hypermethylated. Further, we expect that the larger the difference between the two conditions, the more robust the MSP results will be, when using the predicted primers as biomarkers in a clinical screening context. Therefore, we implemented a filter, which removes from the `bedmethyl` file all positions with a methylation above 10% (default of `--minCpGs`) over all reads mapping to this position in at least one control sample, Fig. 9C. This cutoff can be adapted by the user to increase or decrease specificity by setting a lower or higher methylation threshold, respectively, in control samples. Adaption might be preferable if (i) the input data quality changes or (ii) different PCR assays are used, e.g., when using additional TaqMan probing.

#### Calculation of boxplot data

Next, for all remaining positions, the methylation values for the disease and control samples are aggregated per position and visualized as boxplots (Fig. 9D-G). The 25th and 75th (equals upper and lower quartile, respectively) are calculated with the `numpy` function `nanpercentile`, the 50th percentile (equals median) with the function `nanmedian`. NAN values, where no methylation status could be determined for a sample and position were not taken into account in this way. For the calculation of the 0th and 100th percentile we use the interquartile range *iqr* = 75th percentile − 25th percentile. The 0th percentile was calculated as the lowest value above 25th percentile −1.5 · *iqr*. The 100th percentile was calculated as the highest value below 75th percentile +1.5 · *iqr*. The 0th and 100th percentile represent the highest and lowest value, excluding outliers.

#### Filter 2: Selection for differentiating CpGs

We remove all CpGs, which show no strong differences between disease and control sample. Thus, for each position the median disease methylation (50th percentile) has to be above the 100th percentile of the control sample. As a result, for most arbitrary genomic regions, the methylation in disease samples is only moderately elevated, such as in Fig. 9D, and therefore removed, see Fig. 9F. Potential biomarker regions are expected to be hypermethylated over a longer distance in diseased samples, see Fig. 9E and are subjected to the rest of the workflow, see Fig. 9E/G.

#### Filter 3: Selection of MSP primer suitable regions

Regions for the MSP primer design are selected based on a defined (i) primer length (default 18-24 nt by `--minPrimerLength` and `--maxPrimerLength`) and a (ii) minimum number of differentially methylated cytosines (`--minCpGs` default 3). Thus, the genome is screened for regions which have at least three positions being methylated in the disease samples and unmethylated in the control samples within at most 24 nucleotides. The selected potential MSP regions can contain multiple possible primer positions, see Fig. 9G.

#### Score calculation

For all potential primer regions a score is calculated based on the disease sample coverage and disease sample methylation, see Fig. 9H/I. The primer score is calculated as:

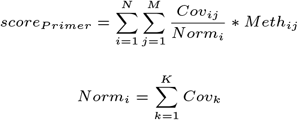

with N = number of disease samples, M = number of CpGs within the primer, and K = number of CpGs within disease sample i. For all remaining CpGs within a primer, the normalized coverage is multiplied by the methylation percentage for each disease sample. In this way, primers with highly covered and highly methylated cytosines in the disease samples result in a high score. As the control methylation has already been checked in Filter 1 and 2, only the disease sample methylation is taken into account. The coverage is normalized to be comparable between all disease samples to receive relative coverage values for each disease sample.

#### Primer pair formation

All potential primer combinations resulting in MSP regions with length between 60 and 400 nt (default, including primers, addressed by `--minAmpliconLength` and `--maxAmpliconLength`) are extracted, with the score being the sum of the two corresponding primer scores, see Fig. 9J/K. Overlapping primer pairs are possible and desired due to a combination of different annealing temperatures and other user-specific requirements.

#### Further output

By default, `diffONT` produces four output files: `boxplotData.tsv` contains aggregated methylation information per position for all differentially methylated cytosines that meet the criteria of Filter 1 and Filter 2. `interestingCpGs.tsv` provides per sample, per position coverage and methylation information extracted from the bedmethyl file for all differentially methylated cytosines that fulfill the criteria of Filter 1 and Filter 2. `primerData.tsv` lists information for each predicted MSP primer. `pcrProduct.tsv` includes details for every predicted MSP region. If the program is run multiple times, these files will be overwritten unless a different output folder is specified. As further output, the DNA sequence for the primer and MSP regions can be reported, if a genome reference sequence is given as input (with parameter `--reference`). For the primers, the DNA sequence is reported in an additional column in the output file `primerData.tsv`. For the MSP regions the DNA sequences are stored in a fasta file `mspRegions.fa`. Additionally, annotated genes overlapping the MSP region are reported in an additional column within the MSP region output file `pcrProduct.tsv`, if an Ensembl (38) genome annotation is given by the parameter `--annotation` in `gtf` or `gff` file format.

This approach is particularly well-suited for analyzing very low coverage data and small sample sizes, as it does not rely on traditional test statistics. While it’s important to acknowledge that lower coverage or fewer replicates may result in a higher likelihood of false positives and false negatives, the method remains robust under these conditions. Additionally, with fewer control samples, the runtime may increase slightly due to the retention of more CpGs for analysis.

Often, multiple potential primer regions are located within close proximity. The `diffONT` github repository https://github.com/rnajena/diffONT/ has the option with a separate script `groupPCRproducts.py` to merge overlapping regions into large regions. These larger regions can then be used for primer design with other tools (e.g., Thermo Fisher’s `Methyl Primer Express` (39)).

ONT benchmarking dataset for comparison with `metilene` and `DSS`

The publicly available Oxford Nanopore Technologies (ONT) Benchmark Dataset rrms 2022.07 was accessed in November 2023 from ONT^3^. In the following, we refer to this dataset as “ONT dataset”. The corresponding human reference genome hg38 (GCA 000001405.15) and annotation (GCA 000001405.15) were retrieved from the NCBI (40) ftp server. The dataset consists of DNA extracted from COLO829 melanoma fibroblasts (ATCC CRL-1974) and COLO829BL normal B lymphoblasts (ATCC CRL-1980) cell-lines, which were established from a 45-year-old male individual with metastatic malignant melanoma. The DNA was sequenced on multiple flow cells per sample, using adaptive sampling to enrich for CpG islands in the human genome, see Tab. 7. Methylation-specific basecalling has been performed, resulting in bam files containing per CpG and strand orientation methylation information (41). We calculated the retained coverage per sample using `mosdepth` (v0.3.3) (42). We used the technical replicates originating from the different flow cells as biological replicates for the comparison of the different tools.

**Table 7.** Benchmarking ONT dataset (43) consisting of five technical replicates for normal B lymphoblasts (COLO829BL) and melanoma fibroblast (COLO829) cell lines. Each technical replicate was Nanopore sequenced on a different flow cell (FC), resulting in an average coverage of about 7X per replicate. The coverage was calculated using the whole genome coverage calculation tool `mosdepth` (42).

|  | FC 1 | FC 2 | FC 3 | FC 4 | FC 5 |
| --- | --- | --- | --- | --- | --- |
| COLO829BL | 7.69X | 6.41X | 6.70X | 6.54X | 7.09X |
| COLO829 | 7.99X | 7.70X | 5.76X | 7.40X | 7.75X |

As input data for `diffONT` we prepared the benchmark dataset using the `modbam2bed` package (v0.9.4) (44). We extracted the methylation information from the ONT output files, which are in bam file format, into bedmethyl-formatted files (45), see STab. S1.

### GIAB benchmarking dataset for comparison with pycoMeth

We could not use the same benchmarking dataset for `pycoMeth` as for the other tools because `pycoMeth` relies on methylation calls from `nanopolish` (9). Re-basecalling and methylation calling with `nanopolish` required the original `fast5` raw data, which was corrupted in the ONT benchmarking dataset, rendering it unusable.

Therefore, for the comparison of `diffONT` with `pycoMeth`, we used the benchmarking dataset from the `pycoMeth` publication (12). However, this GIAB dataset does not contain cancer-specific data. As a result, the comparison with `metilene` and `DSS` was conducted using the ONT benchmarking dataset, which includes the relevant cancer-specific data. We downloaded the samples HG003 (46) and HG004 (47) from the Ashkenazim Trio from the GIAB consortium (48). These samples are from a healthy human male and female, respectively. Basecalling of the `fast5` raw data was performed with `Guppy` (v6.5.7)^4^, using the model `dna r9.4.1 450bps modbases 5mc cg hac.cfg`; methylation calling was performed with `nanopolish` (v0.14.0) (9). For MSP region prediction with `diffONT`, the `bam` output files generated by `Guppy` were merged, sorted and indexed with `SAMtools` (v1.16.1) (49) and converted into `bedmethyl` file format using `modbam2bed` (v0.9.4)^5^, before MSP regions were predicted using `diffONT`. `pycoMeth` DMR detection was performed on the GIAB dataset with default parameters, by running first `CpG Aggregate`, then segmenting the genome into intervals based on methylation by using `Meth Seg` and finally using `Meth Comp` of `pycoMeth` (12) to check for differential methylation for each interval.

### Comparison with tools metilene and DSS (DMR)

For comparison, the ONT dataset was analyzed with `metilene` (v0.2-8) (11), `DSS` (v2.44.0) (14), and `diffONT`. The tool `metilene`, originally designed for analyzing WGBS and RRBS data, requires DNA methylation information in bedgraph format. An in-house bash script converted the data from `bedmethyl` into `bedgraph` file format. In this step, all lines with `NAN` entries in the bedmethyl file (containing no methylation information) were removed. `metilene` was run using the default parameter.

For runtime comparison of `diffONT` with `metilene`, `DSS` and `pycoMeth`, all tools were run on a 64 core processor with CPUs of model AMD Opteron(tm) Processor 6376. All tools were run with default settings for number of threads, if not stated otherwise.

The overlap of predicted regions reported by `diffONT` and `metilene` was determined by using the intersect method from `bedtools` (v2.31.1) (50) with the parameter `-u` and calculating the number of resulting entries. This step was made in both directions, determining the number of `metilene` DMRs overlapping any region predicted by `diffONT` and vice versa.

`DSS` is originally developed for the analysis of WGBS and RRBS data. `DSS` was run in R (v4.2.1) (51) using the package DSS (v2.44.0) (14) with default parameters, requiring a minimum length of 50 nt and a minimum number of three CpG sites per DMR, and 50 % of CpG sites in the

DMR having p-values below 10^−5^. The overlap of predicted regions on chromosome 17 reported by `diffONT`, `metilene` and `DSS` was determined by using the intersect method from `bedtools` (v2.31.1) (50) with the parameter `-u` and calculating the number of resulting entries. The overlap was calculated in all three directions.

### Visualization and CpG island annotation

The Integrative Genomics Viewer (`igv`, version snapshot 05/04/2023) (52) has been used to visualize the read methylation status for the predicted MSP regions. CpG island annotations for the human genome hg38, which are shown in Fig 9, Fig. 5, and SFig. S4C were downloaded from the UCSC genome browser^6^. All plots shown were created using custom-made scripts run under `R` (v4.2.1) (51) using the packages `ggplot2` (v3.5.1) (53), `ggvenn` (v0.1.10) (54) and `dplyr` (1.1.4) (55), unless described otherwise.

## Declarations

### Availability of data and materials

The code of `diffONT` is available at github (https://github.com/rnajena/diffONT/) under MIT license. Data used for the comparison as well as an archived version of the github repository is available under Zenodo.org under the DOI 10.5281/zenodo.14501611.

### Authors’ contributions

DM, EB, LW and MM have made substantial contributions to the conception and design of the work. All authors read and approved the final manuscript.

## Acknowledgments

The authors thank Alfred Hansel and Martina Schmitz for helpful comments and feedback throughout the project.

## Supplement

**Table S1.** The bedmethyl file format contains summarized methylation information extracted from bam files. The bedMethyl file is a bed9+2 file with the following information in its columns: 1) reference chromosome, 2) Start position in chromosome, 3) End position in chromosome, 4) Name of item, 5) Score from 0-1000. Capped number of reads, 6) Strandedness, plus (+), minus (-), or unknown (.), 7) Start of where display should be thick (start codon) – data not shown, 8) End of where display should be thick (stop codon) – data not shown, 9) Color value (RGB), 10) Coverage, or number of reads, 11) Percentage of reads that show methylation at this position in the genome.

| 1 | 2 | 3 | 4 | 5 | 6 | 9 | 10 | 11 |
| --- | --- | --- | --- | --- | --- | --- | --- | --- |
| chr3 | 194688040 | 194688041 | 5mC | 666 | + | 0,0,0 | 12 | 87.50 |
| chr3 | 194688048 | 194688049 | 5mC | 833 | + | 0,0,0 | 12 | 50.00 |
| chr3 | 194688053 | 194688054 | 5mC | 750 | + | 0,0,0 | 12 | 44.44 |
| chr3 | 194688061 | 194688062 | 5mC | 583 | + | 0,0,0 | 12 | 85.71 |
| chr3 | 194688064 | 194688065 | 5mC | 666 | + | 0,0,0 | 12 | 87.50 |
| chr3 | 194688067 | 194688068 | 5mC | 666 | + | 0,0,0 | 12 | 75.00 |
| chr3 | 194688079 | 194688080 | 5mC | 500 | + | 0,0,0 | 12 | 50.00 |
| chr3 | 194688090 | 194688091 | 5mC | 750 | + | 0,0,0 | 12 | 77.78 |
| chr3 | 194688105 | 194688106 | 5mC | 583 | + | 0,0,0 | 12 | 85.71 |
| chr3 | 194688107 | 194688108 | 5mC | 750 | + | 0,0,0 | 12 | 77.78 |
| chr3 | 194688109 | 194688110 | 5mC | 666 | + | 0,0,0 | 12 | 87.50 |
| chr3 | 194688112 | 194688113 | 5mC | 666 | + | 0,0,0 | 12 | 87.50 |
| chr3 | 194688115 | 194688116 | 5mC | 666 | + | 0,0,0 | 12 | 75.00 |
| chr3 | 194688118 | 194688119 | 5mC | 666 | + | 0,0,0 | 12 | 87.50 |
| chr3 | 194688126 | 194688127 | 5mC | 750 | + | 0,0,0 | 12 | 66.67 |
| chr3 | 194688140 | 194688141 | 5mC | 666 | + | 0,0,0 | 12 | 37.50 |
| chr3 | 194688148 | 194688149 | 5mC | 916 | + | 0,0,0 | 12 | 63.64 |

**Fig. S1.**
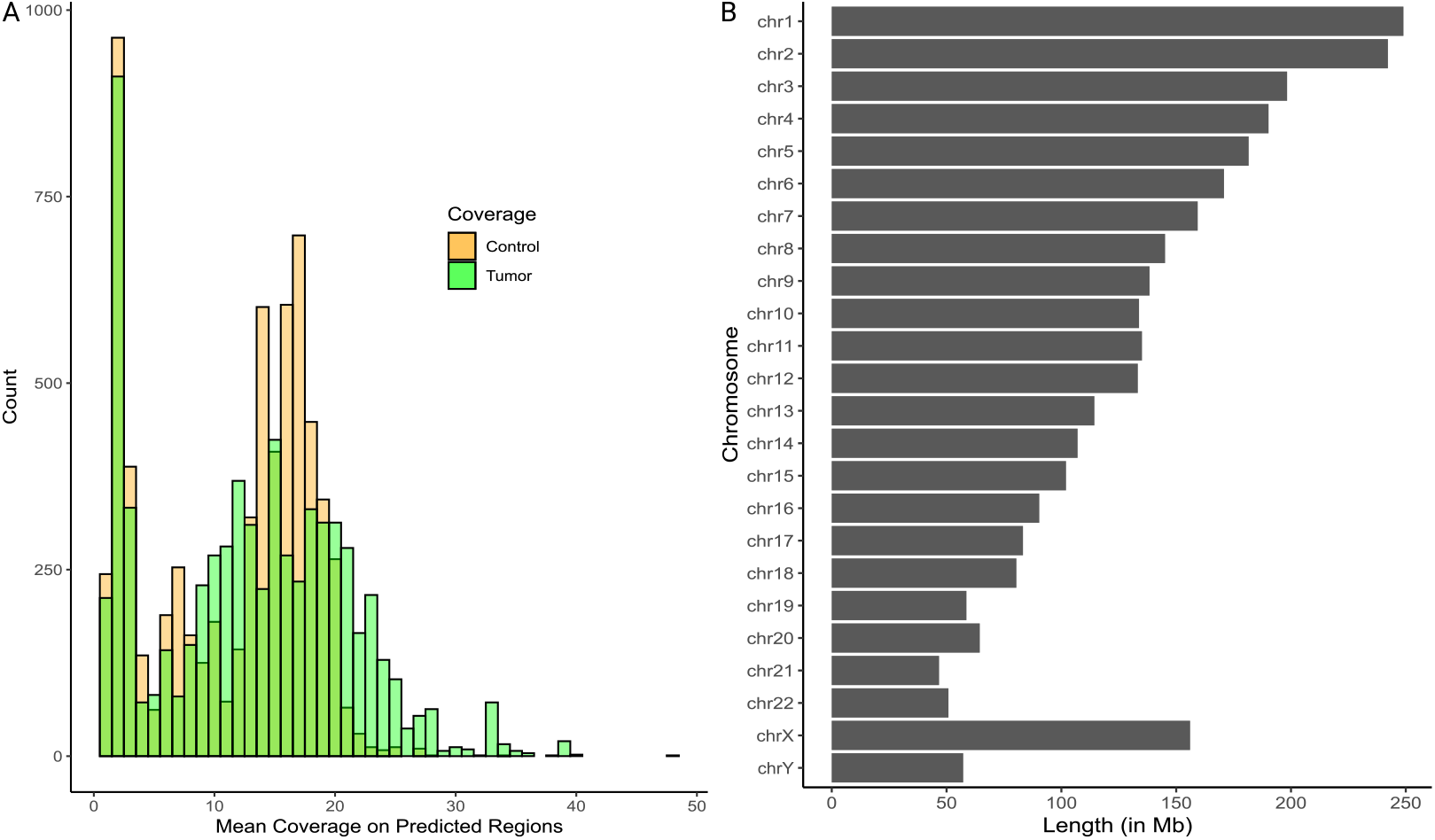
(A) Histogram of the coverage distribution for all regions predicted by `diffONT` on the ONT dataset. Data was binned for coverage, mean coverage per region of control samples and tumor samples is shown in orange and green, respectively. (B) Barplot showing the size of the different chromosomes, based on the hg38 human reference genome. Chromosome length decreases from chromosome 1 to chromosome 22. The size of chromosome X is increased compared to chromosome Y.

**Table S2.** Sequencing coverage per chromosome per flowcell, calculated with mosdepth (42). Sequencing coverage shows differences between the chromosomes over all samples. Especially chromosome Y but also chromosome X show a low coverage (Coverage below 1X for all COLO829 flowcells and below 2X for all COLO829BL samples for chromosome Y). On the other hand, chromosomes 19 and 22 show the highest overall coverage with a mean coverage of 16.76 X on chromosome 19 and 12.89 X on chromosome 22.

| Chr | COLO829 |  |  |  |  | COLO829BL |  |  |  |  |
| --- | --- | --- | --- | --- | --- | --- | --- | --- | --- | --- |
|  | FC 1 | FC 2 | FC 3 | FC 4 | FC 5 | FC 1 | FC 2 | FC 3 | FC 4 | FC 5 |
| chr1 | 8.07 | 7.81 | 5.80 | 7.51 | 7.86 | 8.39 | 6.96 | 7.32 | 7.13 | 7.74 |
| chr2 | 7.29 | 7.04 | 5.31 | 6.76 | 7.12 | 7.11 | 5.95 | 6.20 | 6.06 | 6.58 |
| chr3 | 8.59 | 8.26 | 6.26 | 7.95 | 8.37 | 6.76 | 5.68 | 5.88 | 5.76 | 6.26 |
| chr4 | 7.13 | 6.82 | 5.19 | 6.54 | 6.95 | 6.27 | 5.30 | 5.47 | 5.33 | 5.80 |
| chr5 | 4.55 | 4.37 | 3.31 | 4.18 | 4.44 | 6.62 | 5.55 | 5.76 | 5.65 | 6.12 |
| chr6 | 6.63 | 6.43 | 4.81 | 6.05 | 6.5 | 6.96 | 5.86 | 6.05 | 5.92 | 6.40 |
| chr7 | 11.59 | 11.16 | 8.35 | 10.72 | 11.21 | 8.30 | 6.93 | 7.25 | 7.06 | 7.65 |
| chr8 | 7.30 | 7.05 | 5.32 | 6.75 | 7.12 | 7.04 | 5.92 | 6.12 | 5.98 | 6.49 |
| chr9 | 9.12 | 8.80 | 6.56 | 8.42 | 8.86 | 7.38 | 6.16 | 6.45 | 6.30 | 6.80 |
| chr10 | 5.86 | 5.64 | 4.20 | 5.39 | 5.65 | 8.29 | 6.92 | 7.24 | 7.06 | 7.66 |
| chr11 | 7.96 | 7.69 | 5.74 | 7.42 | 7.73 | 8.31 | 6.9 | 7.21 | 7.02 | 7.64 |
| chr12 | 8.51 | 8.18 | 6.15 | 7.91 | 8.21 | 8.28 | 6.88 | 7.21 | 7.03 | 7.64 |
| chr13 | 5.85 | 5.62 | 4.27 | 5.36 | 5.73 | 5.44 | 4.6 | 4.78 | 4.64 | 5.05 |
| chr14 | 6.80 | 6.57 | 4.92 | 6.33 | 6.64 | 6.68 | 5.57 | 5.84 | 5.70 | 6.19 |
| chr15 | 5.81 | 5.62 | 4.21 | 5.42 | 5.65 | 6.69 | 5.52 | 5.82 | 5.68 | 6.14 |
| chr16 | 9.62 | 9.20 | 6.82 | 8.94 | 9.26 | 11.06 | 9.14 | 9.69 | 9.44 | 10.21 |
| chr17 | 14.08 | 13.59 | 10.05 | 13.3 | 13.53 | 13.52 | 11.05 | 11.78 | 11.43 | 12.37 |
| chr18 | 4.62 | 4.48 | 3.35 | 4.26 | 4.55 | 6.57 | 5.54 | 5.74 | 5.61 | 6.10 |
| chr19 | 18.96 | 18.22 | 13.45 | 17.84 | 18.17 | 18.19 | 14.8 | 15.84 | 15.40 | 16.73 |
| chr20 | 14.22 | 13.75 | 10.21 | 13.28 | 13.68 | 10.33 | 8.53 | 9.03 | 8.81 | 9.50 |
| chr21 | 11.12 | 10.65 | 7.85 | 10.14 | 10.60 | 10.08 | 8.42 | 8.89 | 8.59 | 9.32 |
| chr22 | 16.53 | 15.91 | 11.69 | 15.51 | 15.83 | 12.01 | 9.84 | 10.44 | 10.12 | 10.97 |
| chrX | 4.28 | 4.11 | 3.11 | 3.93 | 4.17 | 3.20 | 2.70 | 2.79 | 2.74 | 2.97 |
| chrY | 0.40 | 0.38 | 0.27 | 0.34 | 0.38 | 1.99 | 1.69 | 1.75 | 1.69 | 1.85 |

**Fig. S2.**
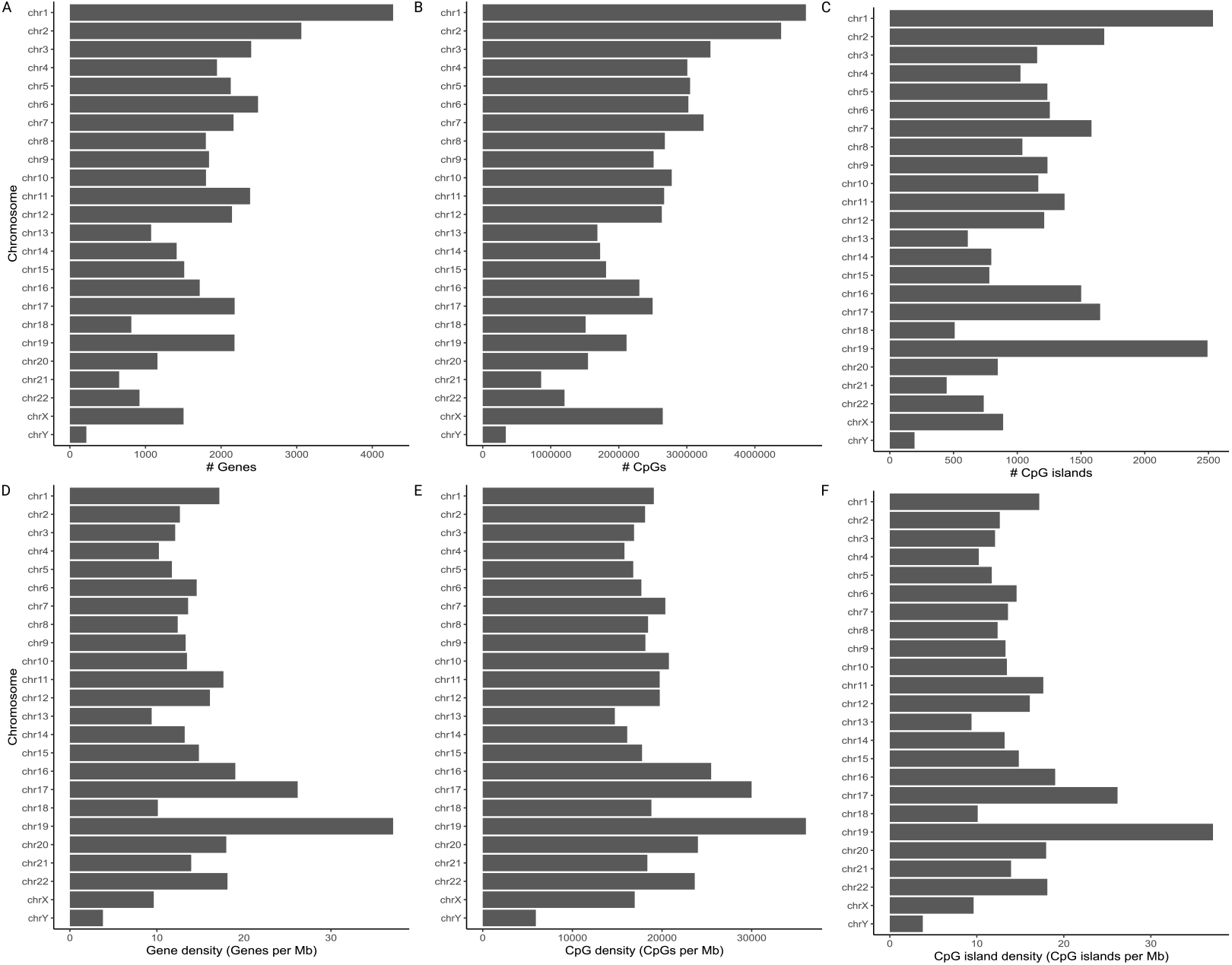
Histogram of the absolute number of (A) Genes, (B) CpGs and (C) CpG islands among the hg38 human genome per chromosome, and (D)-(F) the corresponding densities in relation to chromosome length in [Mb]. Data is based on the human genome version hg38 (GCA 000001405.15).

**Table S3.** Result statistics for the ten highest scored MSP regions predicted by diffONT.

| chrom. | strand | start_fw | end_rev | length | score | cov contr | cov tumor | meth contr | meth tumor |
| --- | --- | --- | --- | --- | --- | --- | --- | --- | --- |
| chr7 | + | 129169295 | 129169493 | 198 | 8978.00 | 15.18 | 34.24 | 0.30 | 97.57 |
| chr7 | - | 128031667 | 128031777 | 110 | 8944.89 | 12.95 | 33.40 | 0.00 | 87.62 |
| chr7 | + | 129169295 | 129169476 | 181 | 8902.96 | 15.15 | 34.20 | 0.30 | 96.46 |
| chr7 | - | 128031397 | 128031690 | 293 | 8684.53 | 13.45 | 33.00 | 0.00 | 85.26 |
| chr7 | - | 128031667 | 128031740 | 73 | 8104.85 | 13.00 | 33.40 | 0.26 | 77.07 |
| chr7 | - | 128031623 | 128031690 | 67 | 8042.99 | 13.02 | 33.30 | 0.00 | 87.00 |
| chr7 | - | 128031669 | 128031777 | 108 | 8028.89 | 12.90 | 33.40 | 0.00 | 87.68 |
| chr7 | + | 129169297 | 129169493 | 196 | 8020.16 | 15.18 | 34.24 | 0.20 | 97.90 |
| chr7 | + | 129169297 | 129169476 | 179 | 7945.12 | 15.15 | 34.20 | 0.20 | 96.78 |
| chr7 | - | 128031671 | 128031777 | 106 | 7940.77 | 12.90 | 33.40 | 0.00 | 86.86 |

**Fig. S3.**
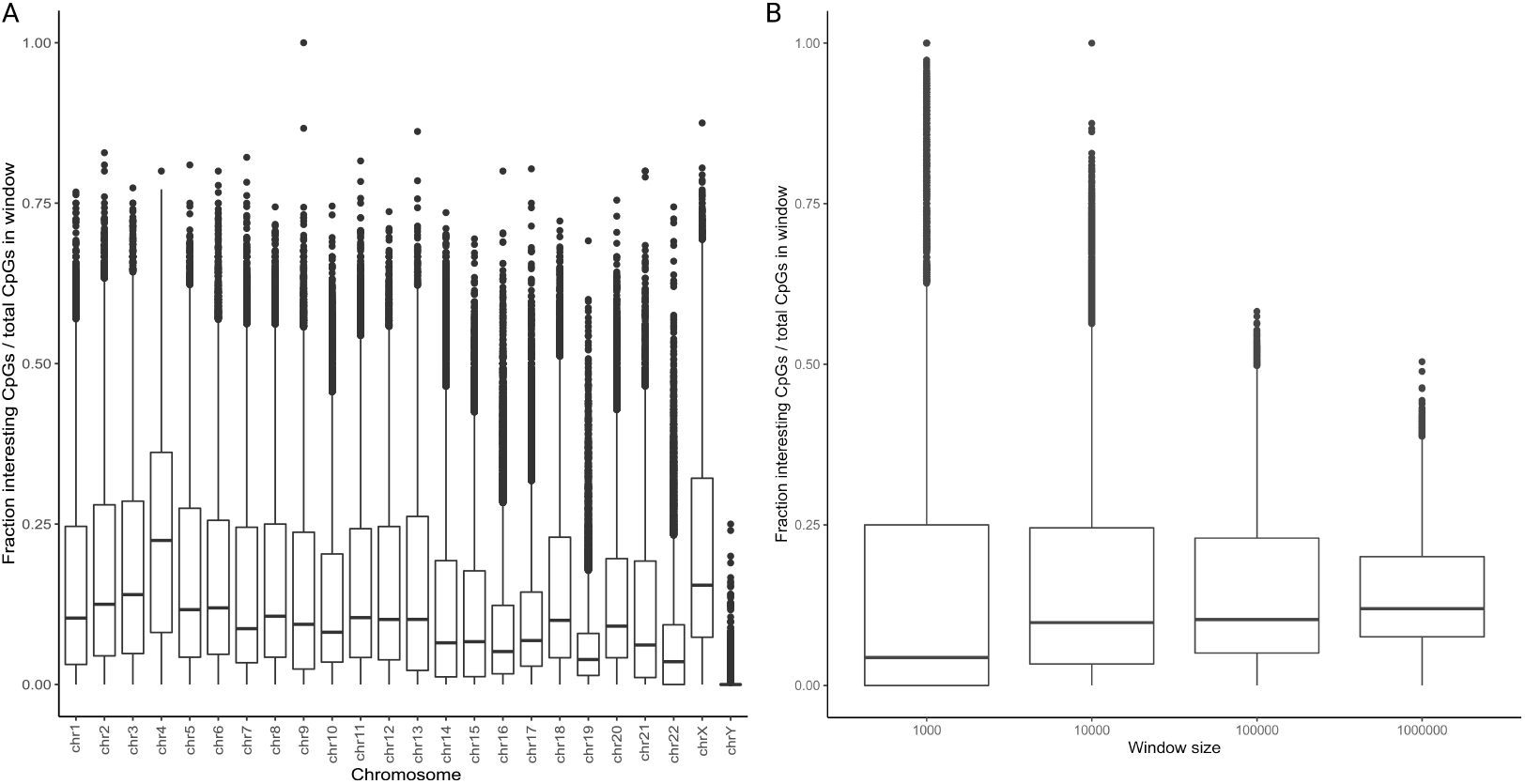
(A) Boxplot showing the ratio of CpGs of interest to total CpGs per chromosome. The fraction was calculated window-wise (size 10 000 nt), which were shifted by half the window length through the genome. The highest median fraction of CpGs of interest is observed on chromosome 4. (B) Boxplot showing the ratio of CpGs of interest to total CpGs within the entire human genome, by using a sliding window approach. CpGs of interest are those CpGs which are passing filter 1 and filter 2. The ratio was calculated for windows of the sizes 1 000, 10 000, 100 000 and 1 000 000 nt. Windows were shifted by half the respective window length through the genome. Larger window sizes result in higher median proportion of CpGs of interest.

**Fig. S4.**
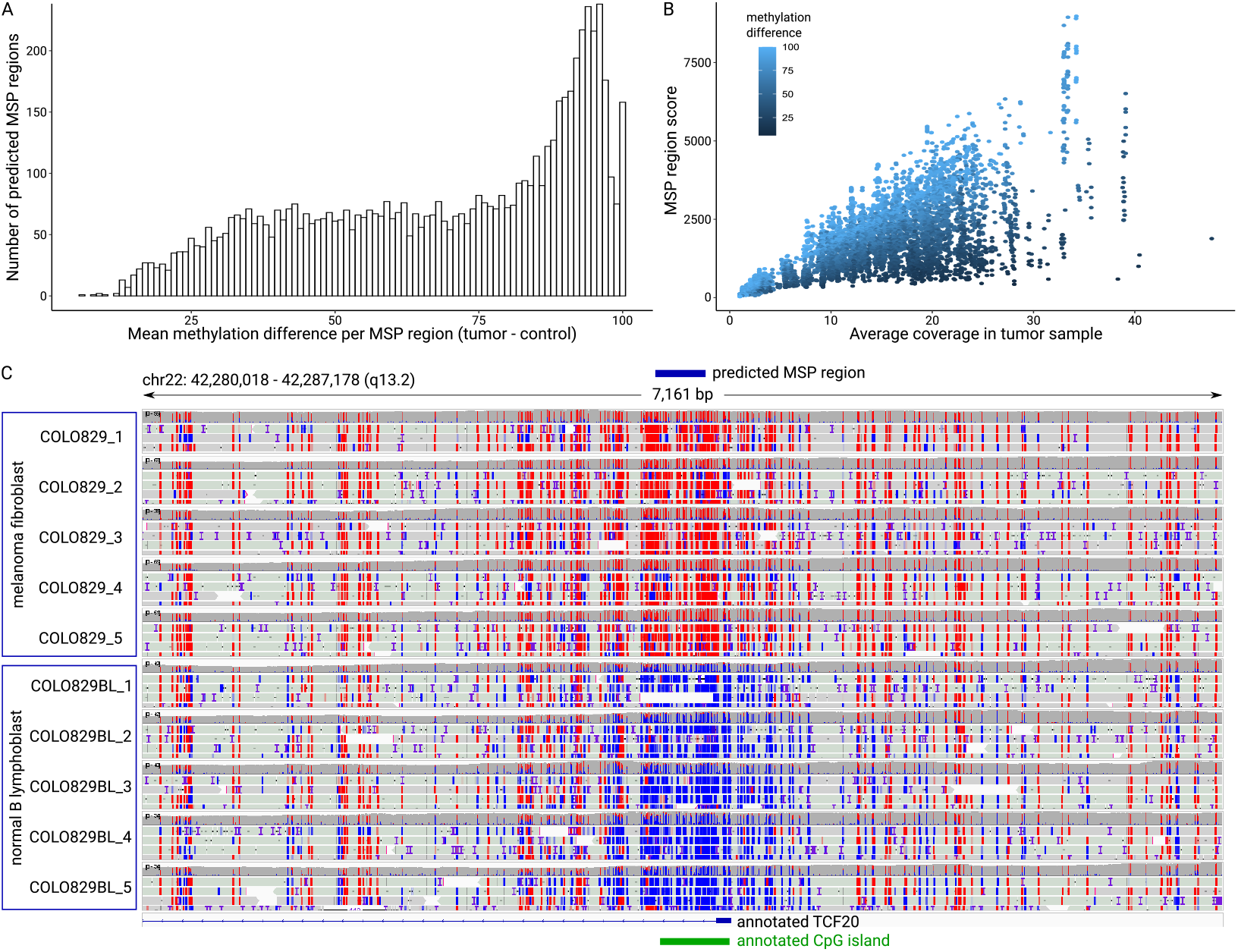
(A) Histogram of the methylation difference values of the regions predicted by `diffONT` on the ONT dataset. The methylation difference is calculated by subtracting the average melanoma fibroblast samples’ methylation from the average normal B lymphoblast samples’ methylation. (B) Visualization of the MSP region score and average methylation in tumor samples for all MSP regions, predicted by `diffONT` on the ONT dataset. The coverage per region shows a gap between 5-8X coverage for the diseased samples. (C) Visualization of one of the top 50 reported MSP regions predicted by `diffONT` on the ONT dataset using the Integrative Genomics Viewer (IGV). At the top the region predicted by `diffONT` is annotated in blue; at the bottom the gene TCF20 is annotated in blue, and an annotated CpG island in green. The 5mC methylation status is color-coded, red = methylated, blue = unmethylated. The melanoma fibroblast sample is shown on top (5x), the normal B lymphoblast sample at the bottom (5x). The predicted MSP region shows strong methylation differences overlapping the TCF20 5’ UTR and promoter region and CpG island.

## Footnotes

1 https://github.com/nanoporetech/dorado

2 https://github.com/rnajena/diffONT/

3 https://registry.opendata.aws/ont-open-data, accessed via `aws s3 ls --no-sign-request, s3://ont-open-data/rrms_2022.07/`

4 Nanopore Community. https://nanoporetech.com/community.

5 https://github.com/epi2me-labs/modbam2bed

6 https://genome.ucsc.edu/cgi-bin/hgTables, accessed in June 2021

